# Trait diversity enhances biomass gains via canopy packing in old-growth but not in disturbed Amazon forests

**DOI:** 10.64898/2026.02.16.706172

**Authors:** Erica Rievrs Borges, Maxime Réjou-Méchain, Grégoire Vincent, Isabelle Maréchaux, Philippe Verley, Jinliu Yang, Ariane Mirabel, Raphaël Pélissier

**Affiliations:** AMAP, Univ Montpellier, CIRAD, CNRS, INRAE, IRD, Montpellier, France; UMR EcoFoG (Agroparistech, CNRS, Cirad, Université des Antilles, Université de la Guyane), Kourou 97310, French Guiana

**Author notes:** Corresponding author: Erica Rievrs Borges **Email:**. **Author Contributions:** Paste the author contributions here. Conceptualization: ERB, MRM, RP. Methodology: ERB, MRM, GV, PV, JY. Investigation: ERB, MRM, GV, IM. Supervision: RP, MRM. Writing—original draft: ERB, MRM, RP, IM. Writing—review & editing: ERB, MRM, RP, IM, GV, AM. **Competing Interest Statement:** The authors declare that they have no competing interests.

**Keywords:** biomass productivity, canopy structure, LiDAR, Forest recovery, disturbance

## Abstract

The existence of a causal link between biodiversity and forest productivity remains largely unexplored in natural systems, especially in hyper-diverse tropical forests. Canopy packing— greater crown complementarity, resulting in more densely packed canopies—has recently emerged as a key structural pathway through which diversity influences forest functioning, though evidence remains limited and sometimes contradictory.In this study, we used repeated airborne LiDAR acquisitions and long-term field monitoring from a tropical logging experiment in French Guiana to quantify canopy packing using the Shannon evenness of plant area density (PAD) and assess its role in mediating the relationship between trait diversity and biomass gains in old-growth and disturbed Amazonian forest stands.Our results show that, in undisturbed forests, functionally diverse communities promote greater canopy packing, which in turn enhances biomass gains. However, this effect was absent in previously logged stands, where forest structural diversity did not fully recover even after 40 years. Our findings indicate that logging reduces canopy structural complexity and disrupts the link between species composition, canopy packing, and productivity in these hyper-diverse, hyper-productive ecosystems.

**Significance Statement:** In this study, measurements from repeated airborne LiDAR acquisitions and long-term field monitoring from a tropical logging experiment in the Amazon forest are used to understand the causal link between biodiversity and forest productivity. The study shows that greater crown complementarity mediates diversity-productivity relationships, with functionally diverse communities promoting greater canopy packing, which in turn enhances biomass gains. However, this effect is lost in disturbed forests. These findings are relevant for understand the ecological mechanisms driving forest productivity and tropical forests response to disturbance and for forest carbon management strategies.

## Introduction

The relationship between biodiversity and ecosystem functions (BEF) remains a central yet debated topic in ecological research, partly because empirical evidence is often inconsistent. A prevailing hypothesis posits that greater diversity enhances ecosystem functioning through niche complementarity, whereby species differ in their ecological strategies and collectively use limiting resources more efficiently (1–3). However, the causal mechanisms underlying these relationships are still largely unexplored in diverse natural systems, particularly in tropical forests. Links between forest diversity and productivity have been attributed in part to canopy structural features such as canopy packing, which can act as a structural mediator arising from greater crown complementary (4, 5). Yet this mechanism remains poorly investigated in natural forests, largely because it is challenging to jointly characterize canopy structure, functional composition, and forest dynamics at spatial scales relevant to ecosystem functioning. Moreover, disturbance history may further obscure or modify BEF relationships (6–8), adding another layer of complexity that may contribute to inconsistent empirical findings (9–12).

Under the niche complementarity hypothesis, tree species diversity is expected to foster canopy packing through a higher functional and structural complementarity among neighbouring trees. Specifically, greater variation in tree shade-tolerance strategies underpinned by contrasting leaf and wood economics traits (13) should lead to vertically structured canopies in which species occupy distinct light niches (14, 15). Additionally, physic based theories predict that the coexistence of contrasted forms, such as leaf sizes or crown architecture, lead to increase canopy space filling (16). Integrating functional trait information is thus critical for interpreting canopy structural variation and for elucidating the mechanisms through which diversity may influence canopy packing.

Most experimental or simulation studies support the idea that greater canopy packing and the associated broader vertical distribution of foliage increased the light-use efficiency of forest stands (17–19). For instance, a recent experiment (20) found that structurally complex stands were twice as productive as simpler ones, particularly when light interception was high, underscoring canopy packing as a major driver of overyielding in diverse forests (3). However, some research suggests that increased asymmetrical competition in structurally complex forests may limit resources per individual, potentially lowering overall ecosystem productivity (10, 21). While the impact of structural complexity on ecosystem functioning in mature forests is not fully resolved, it is even less understood in disturbed forests. Canopy structural complexity is influenced by forest degradation, which is widespread in the tropics (22). For instance, selective logging removes commercial species (often emergent trees) and creates gaps that allow light-demanding species to establish (23). Even though some logging practices aim to maximize forest recovery (24), evidence suggests that canopy structure and species composition need much longer than biomass stocks to recover (10, 23, 25). While old-growth forests typically display multilayered canopies, broad tree-size distributions, and diverse crown architectures, secondary forests tend to have more uniform tree sizes and reduced architectural variation (10, 26), potentially limiting complementarity effects (7, 8) and thereby lowering overall resource-use efficiency (9, 27).

Finally, at least two methodological challenges arise when investigating canopy packing and BEF relationships in forest systems. First, most studies overlook the fact that diversity may influence different biomass components separately or simultaneously through distinct ecological mechanisms (8) potentially blurring interpretations. To address this issue, a recent study analytically disaggregated field-based biomass estimates to separate the structural components—those directly linked to canopy packing (the number of trees and the tree volume)—from the compositional component (wood density; 8). Second, measuring canopy packing independently from the same field data used to estimate biomass is necessary to avoid circularity, but it remains technically challenging: traditional field methods are time-consuming, logistically difficult, costly, and often impractical. Remote sensing, especially LiDAR (Light Detection and Ranging), offers a powerful tool for assessing and mapping forest structure, but below-canopy structural measurements are often biased by scan properties, leading to imprecise estimates. Here again, recent development based on ray tracing approaches have considerably improved unbiased below-canopy structural measurements, offering new opportunities to measure the vertical complexity of forest stands (28).

In this paper, we investigated whether canopy packing mediates BEF relationships in an old-growth and experimentally disturbed Amazon forest located in northern French Guiana and monitored for ca. 40 years. We determined pathways by which tree communities’ trait diversity (functional dispersion) increased canopy packing, characterized through an innovative LiDAR ray-tracing approach that corrects for scan properties (28) at different stages of the post-disturbance forest dynamics. We expected canopy packing to be greater in tree assemblages composed of species with dissimilar resource-use strategies and forms (H1). To measure canopy packing, we used an index derived from the Shannon Evenness Index (see Eq. 2), which measures how evenly is the distribution of plant material (Plant Area Index, PAD) across the forest vertical layers. We also expected that spatially and functionally complementary canopies would lead to denser and more vertically structured canopies, and ultimately, to higher biomass gains (H2; (20)). We determined pathways linking canopy packing to coarse wood productivity (biomass gains), through the net changes in above-ground biomass (AGB) and its structural components (number of trees and above-ground volume) obtained from the analytical disagregation of AGB (Fig. 1). Besides, as the development of the forest vertical structure could lead to density-dependent mortality mediated by shading (6–8), we implemented the same pathways described above for biomass loss, and assessed the relative contribution of biomass gains and losses to total biomass net change. Finally, we hypothesised that stronger canopy packing effects should be found in old-growth forests, as in disturbed forests the shift toward high-light environments favors a narrower set of species able to exploit these conditions and thereby hinders complementarity effects as species dominance becomes the main driver of productivity (H3; (6-8; van der Plas, 2019)).

**Figure 1.**
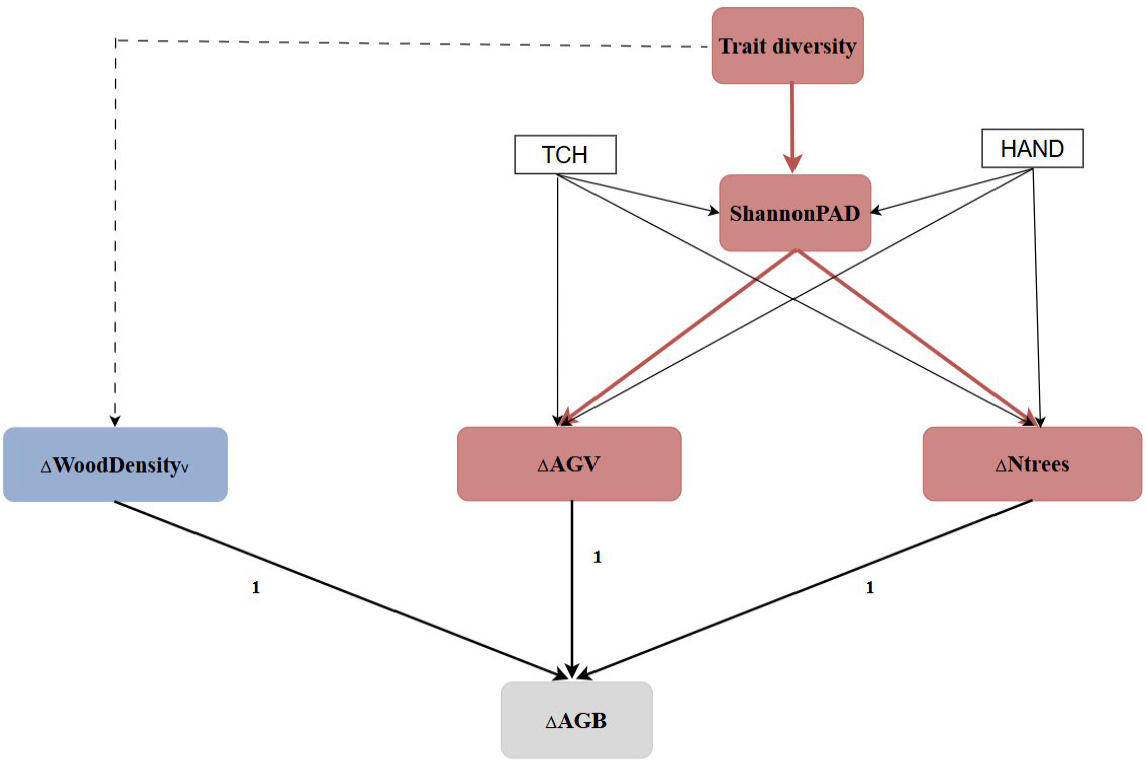
Conceptual model. Description of the expected links among functional trait diversity, canopy packing (vertical structure complexity represented by the Shannon plant area density evenness, ShannonPAD) and ΔAGB components. We considered each biomass component separately, following the approach developed by (29) and applied by (8). In this framework, stand-level above-ground biomass (AGB), defined as the sum of individual tree AGB estimates, is decomposed into the number of trees, mean individual above-ground volume, and mean volume-weighted wood density. After log transformation, this decomposition leads to an additive formulation. Further details are provided in (8). AGV is the log-transformed mean individual above-ground volume, Ntrees, the log-transformed number of trees, AGB, the log-transformed aboveground biomass and Δ represents their net change. As an exact (analytical) relationship, the effect of ΔAGV and ΔNtree on ΔAGB has a known coefficient value of 1 (no estimation is required here due to the analytical solution). We considered two additional covariates in our model: TCH, top-of-canopy height, and HAND, height above the nearest drainage. The wood density component of AGB in blue (dotted line), which is expected to be related to sampling effects rather than canopy packing (8), is not included in our model.

## Results

Canopy packing, estimated here by the LiDAR-derived Shannon evenness index of plant area density (PAD), was statistically lower in forest plots disturbed 40 years ago by both logging and thinning (L2) compared to control old-growth and the logging-only treatment (L1) (Fig. 2 and S1). We specifically found a trend of decreasing Shannon PAD evenness as past logging intensity increased. Accordingly, as exemplified in Fig. 3 (see Fig. S2 for all subplots), the canopy profiles and the respective cross-sections illustrated that the PAD was better distributed across the understory in control plots, while the most intensively logged plots showed a peak in PAD at 20 m and less homogeneous vertical distribution of plant material (Fig. 2 and S2).

**Figure 2.**
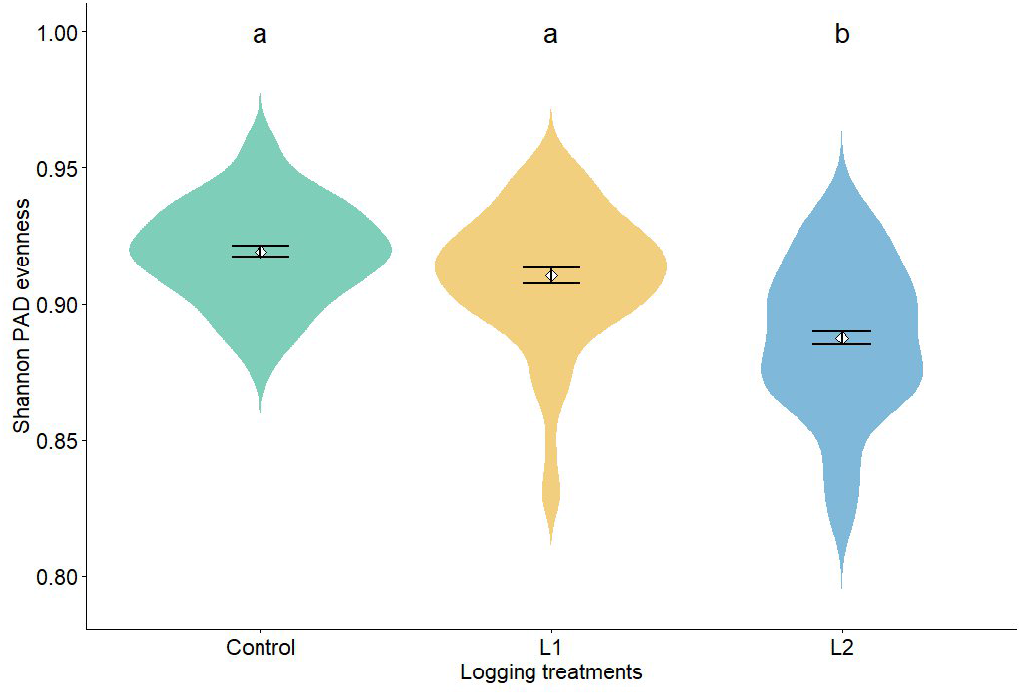
Violin plots showing the distribution of Shannon’s evenness Index of Plant Area Density (PAD) across the logging treatments in 2015. Higher Shannon evenness index of plant area density (PAD) is predicted for Control and L1 plots compared to L2 plots [Tukey HSD test F-value=38.5, p-value<0.001]. “Control” represents undisturbed plots, “L1” represents plots with logging treatment only, and L2 represents plots with both logging and thinning treatment. The width of each violin represents the kernel density of Shannon PAD evenness values. White diamonds indicate mean values, and black error bars represent ±1 standard error of the mean. Different letters (a, b) indicate significant differences in a pairwise comparison of least-squares means (Tukey HSD).

**Figure 3.**
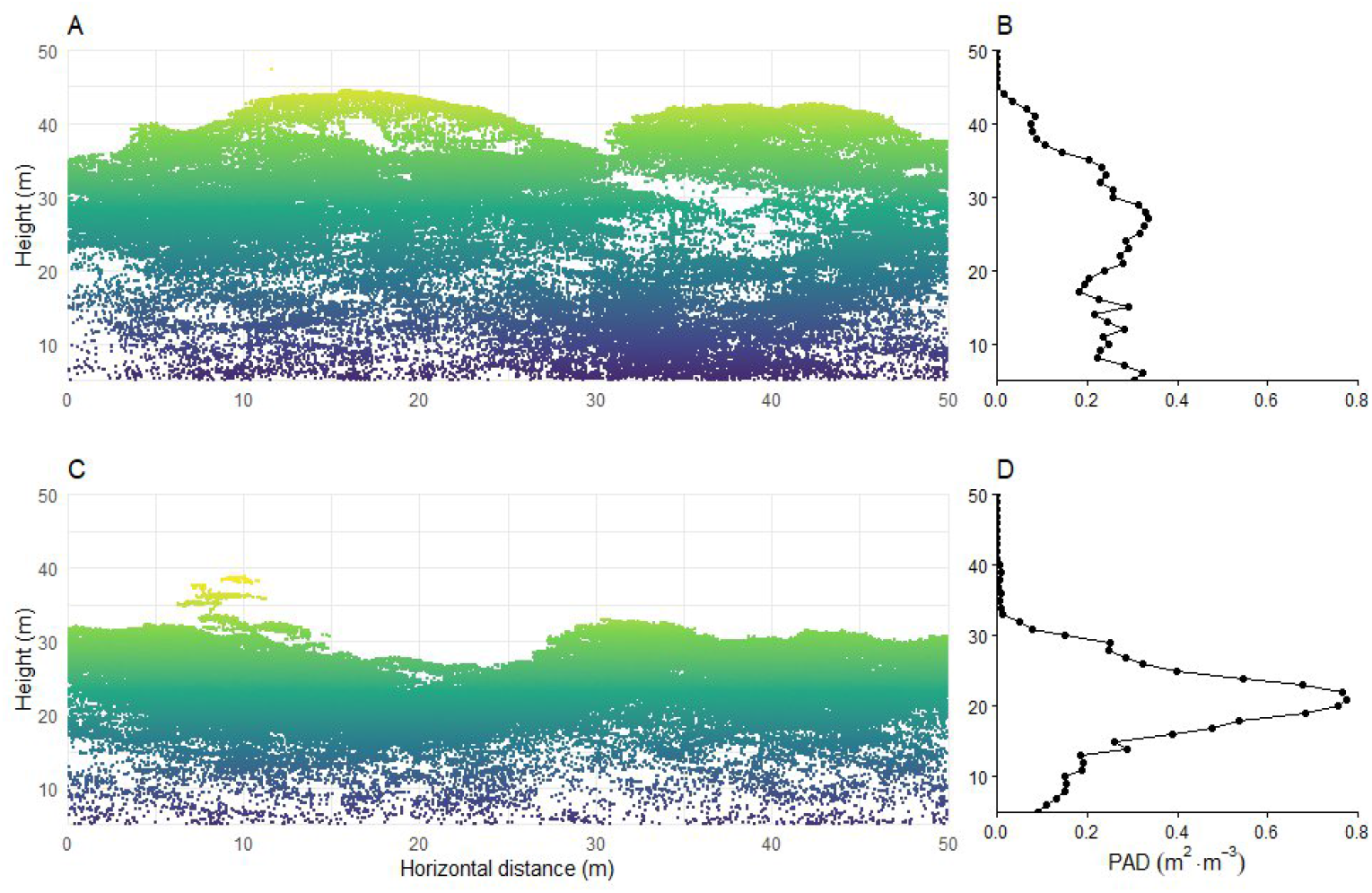
Examples of vertical canopy structure observed in a control and a disturbed 0.25-ha subplot. A and C: LiDAR point cloud coloured according to height above the ground; B and D: variation of Plant Area Density (PAD) means per 1-meter layers across the vertical canopy profile, with PAD distributions modelled from the LiDAR data with the AMAPVox software. A and B: control old-growth treatment; C and D: L2 logging and thinning treatment

Multivariate and trait-by-trait functional dispersion (except for SLA) were statistically greater in L1 plots than in control and L2 plots for the two studied years (Fig. S3). Multivariate functional dispersion had a significant positive effect on Shannon evenness of PAD within old-growth plots (Fig. 4 and S4), even though HAND, a topographic index representing the height above the nearest drainage, was the best predictor of Shannon PAD evenness with more densely packed canopies in the wettest areas (small HAND values) for control and LI plots (Fig. 4 and S4, Table S1 and S2). Trait-by-trait (univariate) analyses revealed that the positive effect of functional dispersion on canopy packing was mostly due to the functional dispersion of leaf area (for 2015, Fig. 4) and leaf thickness (for 2019, Fig. S4). By contrast, the functional dispersion of the other traits showed no significant effect on Shannon PAD evenness (Fig. S5). As past logging intensity increased, the positive and negative effects of functional dispersion and HAND, respectively, decreased with no significant effect of functional dispersion on Shannon PAD evenness in either logging treatment. By contrast, the effect of canopy height (TCH) increased along the logging intensity gradient, with a significant effect in the L2 treatment.

**Figure 4.**
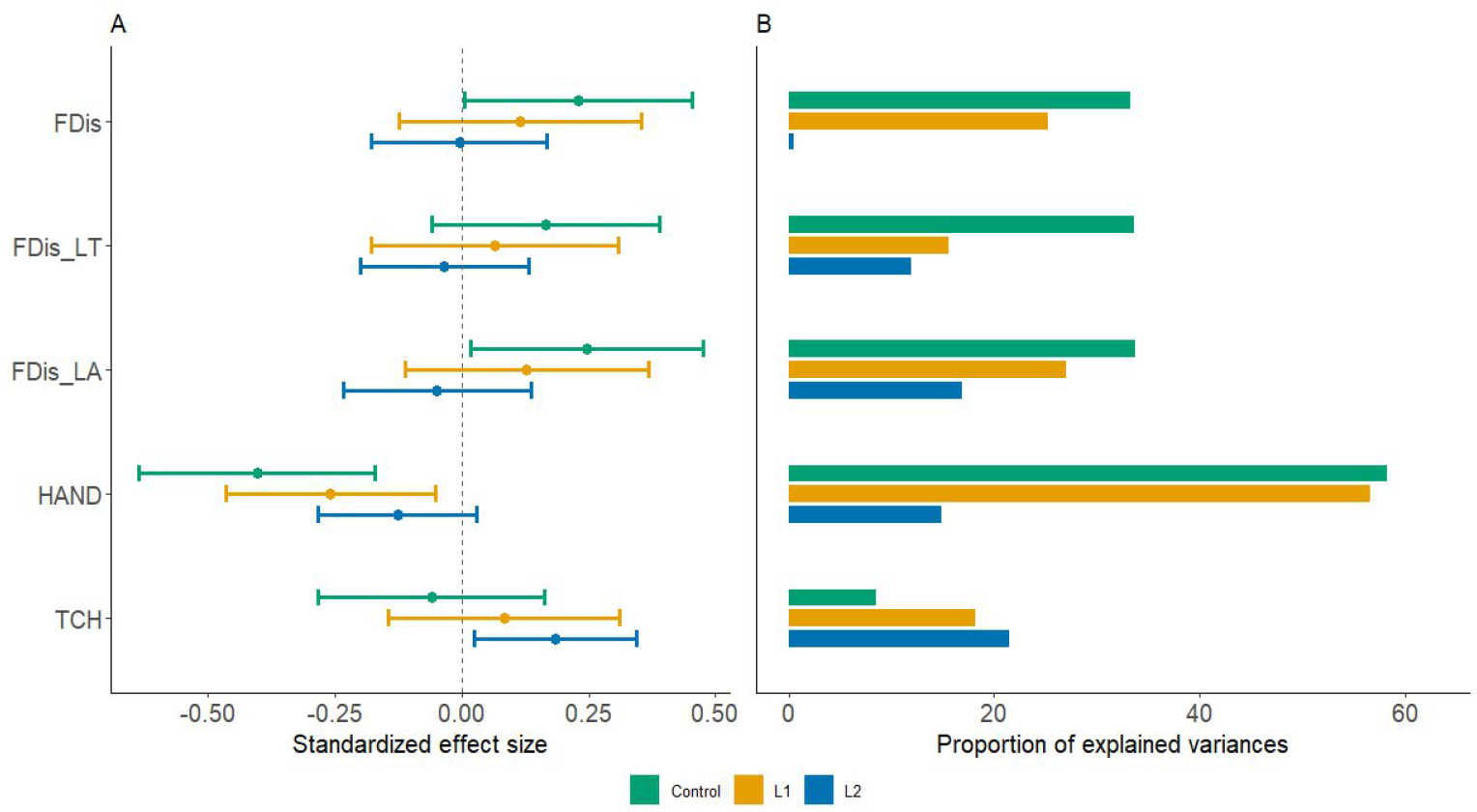
Effects of multiple drivers of canopy packing, estimated using Shannon PAD evenness. Standardised effect sizes (A) and proportion of explained variances (B) of the significant predictors of Shannon PAD evenness for 2015. Control, old-growth; L1, logging only treatment; and, L2, logging and thinning treatment. TCH, top-of-canopy-height; HAND, height above nearest drainage; FDis, functional dispersion; LT, leaf thickness; and, LA, leaf area. Results from the multivariate FDis model are shown, while the univariate LT and LA models are included for context.

To understand the impact of canopy packing on the structural components of biomass gains and losses we assumed that Shannon PAD evenness increases the number of trees and/or their mean above-ground volume, but does not influence the wood density component since the latter is expected to be related to the sampling effect (Fig. 1; 8). We studied two time intervals to calculate biomass gains (2015–2019 and 2019–2023), with the Shannon PAD evenness measured at the beginning of each interval (2015 and 2019, respectively) as predictor of subsequent biomass gains. For biomass gain, Shannon PAD evenness showed a significant positive effect on tree recruitment (ΔNtrees; Fig. 5a, Fig. S6) for both studied intervals and all treatments except L1, while the Shannon PAD evenness effect on ΔAGV was significantly positive for control plots in the first interval but not significant on the other cases (Fig. 5a, Fig. S6). Finally, Shannon PAD evenness did not show an overall influence on biomass loss by tree mortality, (but see Fig. S6 for the 2019-2023 interval). Across all treatments and measurement intervals, biomass gains contributed more strongly to net AGB change than biomass loss, a pattern particularly pronounced in the L2 treatment (Fig. 5, Fig. S6).

**Figure 5.**
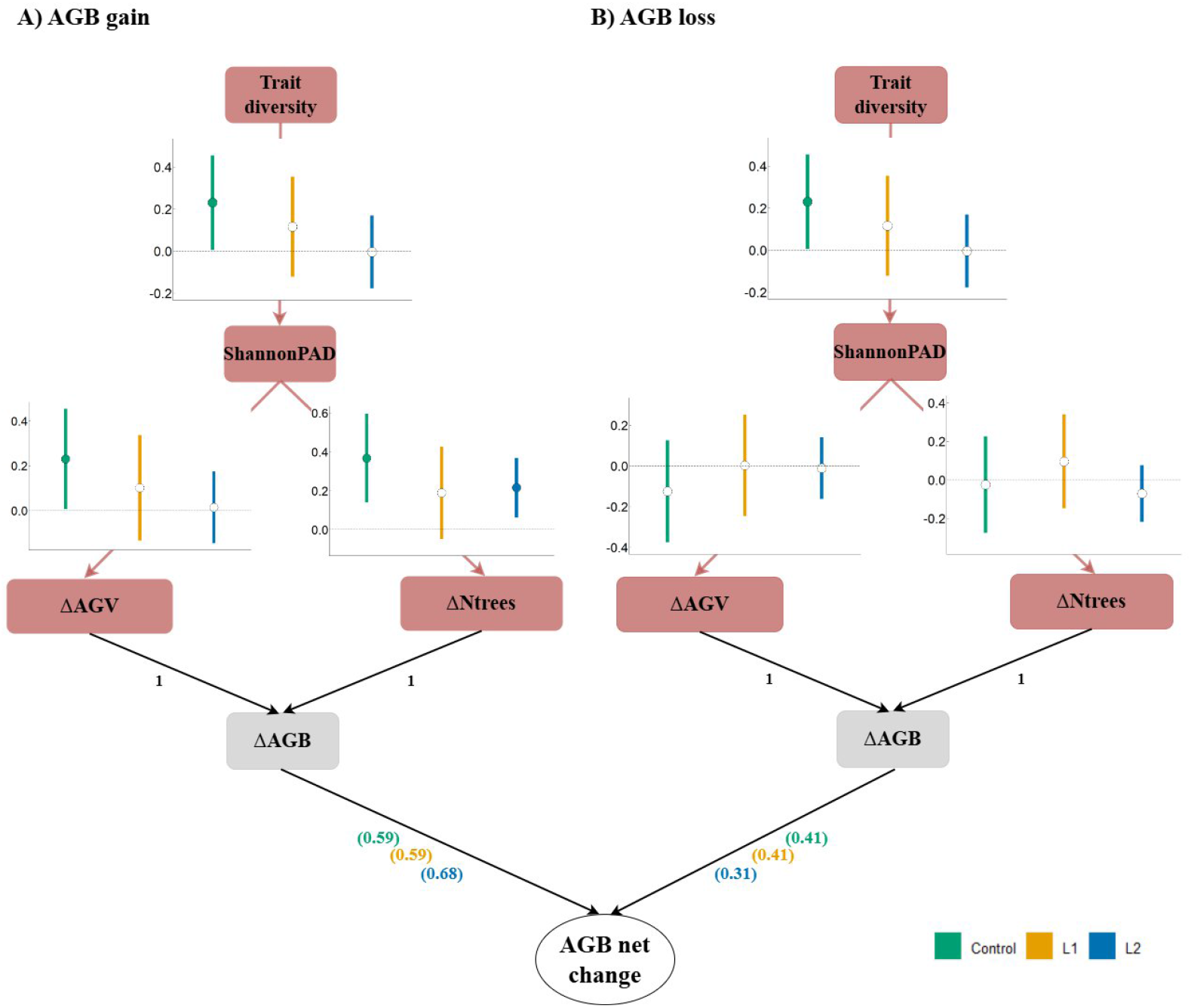
Structural equation models describing multiple drivers of Shannon PAD evenness and biomass gains and loss. Results of the structural equation models for the effect of multivariate functional diversity on structural diversity (Shannon PAD) and the components of above-ground biomass gains (A) and loss (B, ΔAGB) for the first monitoring interval (2015-2019; see Fig. S6 for the 2019-2023 interval). Control, old-growth; L1, logging and; L2, logging and thinning treatment. ΔAGV, change in log-transformed mean individual above-ground volume; ΔNtree, change in log-transformed number of trees. The co-variables HAND and CHM are not shown in the SEM for visualisation clarity (results shown in Tables S1 and S2).

Shannon PAD evenness was the most important driver of biomass gains in control plots, with the highest proportion of explained variance for both intervals (Fig. 6 and S7). However, the same pattern was not found in disturbed plots. Besides Shannon PAD evenness, TCH generally showed significant negative effects on biomass gains (but see the control results for the first interval in Fig. 5), with stronger effects on L1 and L2 (Fig. 6B and S7). The effect of HAND on biomass gains was not significant and inconsistent among treatments and interval years (Fig. 5 and S7).

**Figure 6.**
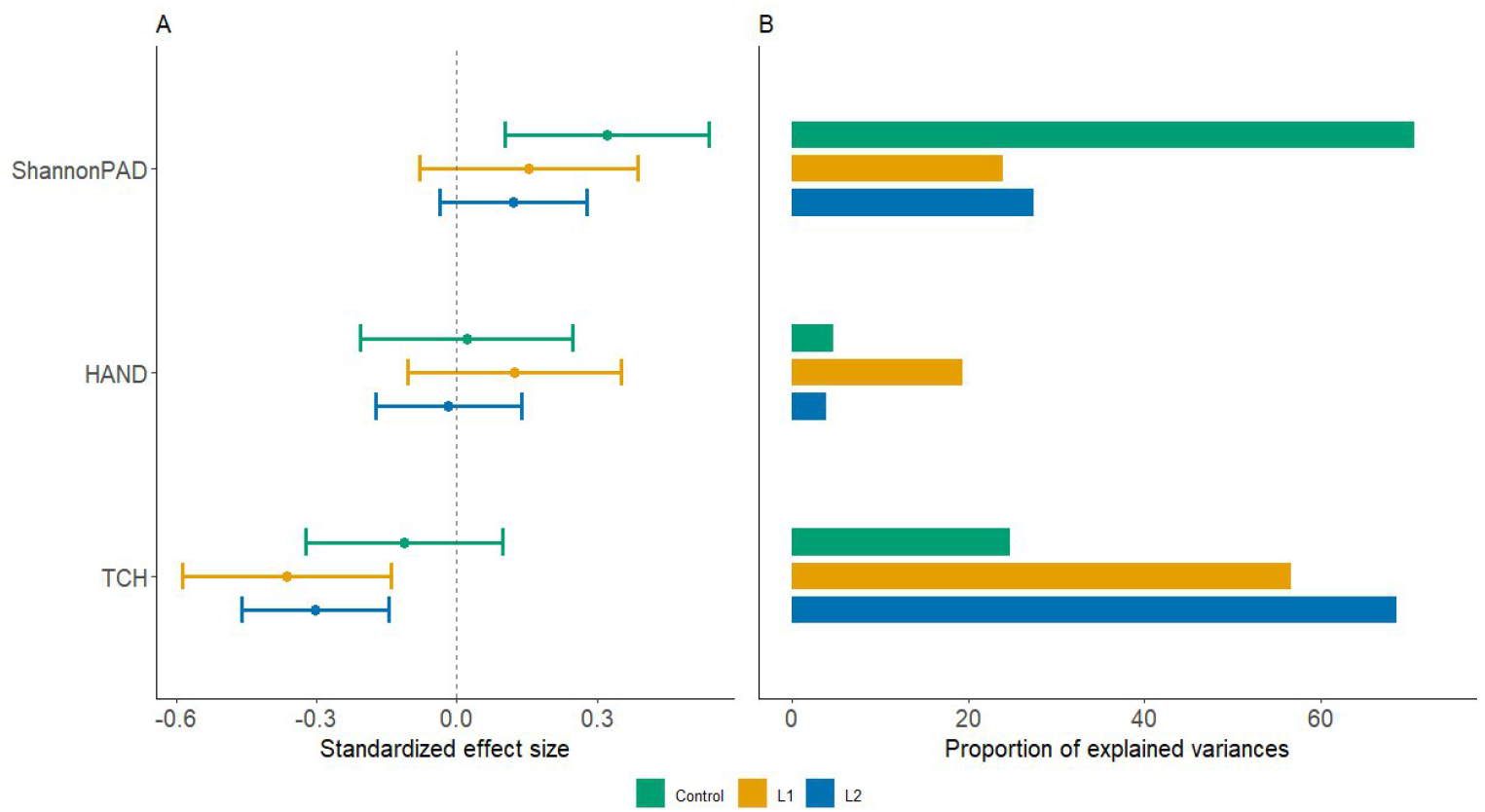
Effects of multiple drivers of biomass gains. Standardised effect sizes (A) and proportion of explained variances (B) of biomass gains predictors for the 2015-2019 interval for the different treatments (see Fig. S7 for the 2019-2023 interval). Control, old-growth; L1, logging and; L2, logging and thinning treatment. HAND, height above nearest drainage, TCH, top-of-canopy-height.

Overall, our models explained between 9% and 17% of the variance in Shannon PAD (fixed effects; Tables S1 and S2). Notably, the relative importance of each predictor varied substantially across treatments. For example, functional dispersion accounted for 7% of the variation in Shannon PAD evenness in L2 plots, but 33% in control plots (Fig. 4B and S3B). Similarly, although the explained variance of biomass gains was comparable across control and disturbed plots (R^2^m = 0.12, 0.14, and 0.10 for control, L1, and L2 for the first interval, respectively), the contribution of Shannon PAD evenness differed markedly: 70% in control plots, versus 24% and 27% in L1 and L2 (Fig. 6). As an illustration, in control plots, increasing multivariate functional dispersion by 30% (roughly equivalent to a two-standard-deviation increase) would lead to a rise in Shannon PAD evenness of around 1%. However, a 5% increase in Shannon PAD evenness (also roughly equivalent to a two standard deviations) produced a stronger effect, enhancing biomass gains by almost 30%. In the second interval, the differences across treatments were less pronounced (Fig. S7).

## Discussion

### Paste your discussion here

At a large-scale logging experimental site, we showed that functional dispersion, particularly traits related to shade tolerance and resource acquisition, increased biomass gains through canopy packing in undisturbed Amazonian plots. However, this effect was absent in previously disturbed stands, where forest structural diversity did not fully recover even ca. 40 years after logging activities subsided, a pattern reported here for the first time in Amazon forests. Our findings indicate that the loss of large trees from the logging treatments drastically changed the ecological processes driving biomass gain, disrupting the functional link between species composition, canopy packing, and biomass gain.

### Functional diversity increases canopy packing in old-growth forests

Confirming our first hypothesis, our results showed that functionally diverse communities enable a better distributed occupation across vertical forest layers (increased canopy packing) in old-growth forest plots. This is driven by differences in key functional traits, reflected in contrasting leaf morphologies. This result also held true for univariate functional dispersion of leaf area and leaf thickness, which together capture key aspects of light-use and carbon investment strategies (13, 30). Leaf thickness varies with canopy position: sun leaves are typically thicker than shade leaves, allowing them to tolerate higher irradiance and reduce water loss, while thinner leaves in shaded environments are adapted to maintain function under low light (13, 31–35). Leaf area also varies among species and influences light interception; a diversity of leaf areas within a community can enhance light use efficiency by optimizing canopy space filling and capturing sun flecks in the understory (36). A diversity in successional strategies foster such trait dispersion by allowing the stratification of light-demanding species that dominate the upper canopy soon after gap formation (and tend to persist for decades) and shade-tolerant species that establish in the understory before some eventually grow into the top of the canopy (30, 37–39). Together, these complementary strategies lead to greater vertical partitioning of space, reduced competition, and enhanced canopy packing (40). Therefore, our findings confirm that higher canopy packing is achieved in communities characterised by higher functional complementarity (represented in this study by functional dispersion), where shade-tolerant species co-exist with light-demanding species, fostering the formation of functionally diverse tree species communities in terms of physiological adaptation to shade (41).

### Canopy packing increases biomass gains in old-growth forests

We found that higher Shannon PAD evenness was associated with greater biomass gains in old-growth forest plots, confirming our second hypothesis. As stated above, a more homogeneous distribution of canopy space is associated with improved niche partitioning, driven by more efficient allocation of space and light resources. In older forest stands, as previously found at the same study site (42), gaps created by large trees falling are more frequent, allowing niches to be refilled by small trees (43, 44). Those small trees do not directly compete with the largest trees due to vertical stratification and their different sizes and shapes, but they can contribute to increased productivity in the whole stand by enabling a more complete use of resources (45).

Indeed, the canopy packing effect was strongly related to tree recruitment (ΔNtree). These findings indicate that niche complementarity (possibly related to gap dynamics) is an important determinant of forest productivity (46, 47).

Notably, Shannon PAD evenness was the most important variable with a substantial explanation of biomass gains in old-growth forest plots, especially for the earliest 2015-2019 interval, surpassing the influence of water availability (HAND). This finding underscores the central role of canopy structural organisation in regulating forest productivity. While water availability is widely recognised as a primary control of tree growth in tropical forests at large scales (48), our results suggest that fine-scale structural complementarity may be equally, or even more, important in light-limited forests. However, in addition to functional dispersion, water availability was also an important driver of canopy packing, possibly indicating that soil moisture and reduced hydraulic stress in wetter topographic positions foster higher understory density (49), thereby contributing to increased canopy packing and, indirectly, biomass gains. Moreover, in our studied plots, bottomland positions were previously found to exhibit higher tree turnover—both mortality and recruitment—than upland sites, with hydrological variability and mechanical stress creating gaps that further promote understory growth and canopy rearrangement (50).

### Functional diversity effect on biomass gains mediated by canopy packing is lost in disturbed forests

Finally, consistent with our third hypothesis, we showed that the effect of functional dispersion on canopy packing is context-dependent, with functional dispersion not increasing Shannon PAD evenness in disturbed plots. Disturbed plots are still recovering in terms of functional composition, as they are more dominated by species with acquisitive traits than mature plots (Fig. S8; see also (51) for more detailed results of functional composition trajectories across all treatments at the genus level), even though their functional composition does not seem to affect the functional dispersion, with L1 plots showing greater functional dispersion than control plots (Fig. S3). In the more open and resource-rich conditions created by disturbances, trees are less constrained by neighbours and can expand or reposition their crowns opportunistically into newly available light and space. This relaxed competition, combined with the altered microenvironment created by the logging legacy, weakens the role of species’ intrinsic light-use strategies in determining their vertical positioning. As a result, functional differences among species become less tightly linked to vertical niche partitioning, which explains why even high functional dispersion in L1 plots does not translate into greater vertical stratification. Instead, in disturbed plots, Shannon PAD evenness was explained by abiotic controls such as water availability (HAND in L1 plots) or stand stature (TCH in L2 plots), in agreement with recent findings that environmental and structural factors can outweigh biotic factors in shaping forest structure (52).

In disturbed plots, Shannon PAD evenness showed no significant effect on biomass gain (even though it positively affected ΔNtree for the most disturbed treatment). Instead, we found that TCH was the most important driver of biomass gains in these plots, with a significant negative effect. This likely reflects smaller-scale successional gradients within treatments, where plots with lower canopy stature—typically representing earlier successional stages (46) —exhibit higher productivity during recovery (Fig. 6 and S7). Our results revealed that the relative influence of canopy packing on biomass gain declined, while the importance of canopy height increased along the disturbance gradient. Canopy height also explained variation in biomass gain in control plots, but this effect was proportionally greater in disturbed plots (∼58%) than in control plots (33%; Table S1, S2). These findings suggest that logging altered the balance between structural and compositional drivers of biomass gain, weakening the role of functional diversity and increasing the influence of canopy stature.

Recent studies have shown that multi-layered forest canopy results in more stable microclimate conditions (53, 54), more homogenous thermal conditions in the understory (55) and might even be more resilient to environmental stressors (56), indicating the importance of measuring and potentially managing forest structural diversity. It was also observed that different management interventions led to shifts in species composition, influencing tree canopy attributes that, in turn, affect ecosystem processes and services (54). Our results extend this perspective by showing that the relationship between diversity, canopy packing, and productivity is strongly context dependent (57, 58), and that, while enhancing biodiversity or canopy complexity may be beneficial in some circumstances, generalizations of nature-based initiatives (57, 58) —especially those aiming to boost biomass productivity—can be misleading, as biomass dynamics in tropical forests are governed by multiple interacting processes and mostly by the different land-use histories of secondary forests. Because one third of tropical forests are secondary (22), the positive effect of diversity on biomass gain is likely limited to only a portion of tropical forests.

Therefore, rather than promoting universal prescriptions, restoration and management strategies should therefore be tailored to the local successional context, explicitly accounting for past disturbance, stand age, and the dominant mechanisms operating at each stage.

## Materials and Methods

### Paste your materials and methods here

If your research involved human or animal participants, please identify the institutional review board and/or licensing committee that approved the experiments. Please also include a brief description of your informed consent procure if your experiments involved human participants.

### Study sites and forest inventory data

We used forest inventory data from 12 6.25-ha (250 × 250 m) permanent plots that belong to a logging experimental site at the Paracou Tropical Forest Research Station (5°18′ N, 52°53′ W) in French Guiana. This station is one of the richest tropical forest sites in terms of the availability of ground-level forest monitoring data (currently 118.75 ha measured, 75 ha since 1984; (57). Mean annual precipitation is 3,041 mm year−1, with a pronounced dry season (<100 mm month−1) from August to November. The topography of the study area is globally gently undulating (between 2 and 40 m asl) but includes some relatively steep local slopes (6-20%). A dense hydrographic network forms a series of shallow depressions and low hills. Forests in Paracou are classified as tropical lowland, terra firme forests. Twelve plots were established from 1984 to 1988. Of these, nine plots were subjected to various sylvicultural treatments at different intensities from 1986 to 1988, and three plots were left untouched as controls. Here, we separated disturbance categories based on total biomass loss from silvicultural treatments. Total original biomass loss per treatment was 12-33% for L1 (logging; 3 plots) and 33-56% for L2 (logging and thinning; 6 plots), even though originally L2 contains two slightly different treatments (T2: logging and thinning; T3: logging and thinning and fuelwood) (57). Separating the T2 and T3 treatments yielded similar results (not shown); therefore, to simplify presentation, we decided to pool them into a single treatment.

All trees with a diameter at breast height (DBH) ≥ 10 cm were individually marked, geo-referenced, botanically identified and re-censused every two years. To ensure the quality of the inventory dataset, we corrected for abnormal annual increases and absolute decreases in DBH by aligning each DBH measurement with the individual’s overall growth trajectory (59). Abnormal decreases in DBH were frequently associated with shifts in the measurement point due to the emergence of trunk irregularities, such as buttresses. Each 6.25-ha plot was gridded into 25 50 × 50 m (0.25 ha) subplots, yielding 75 subplots for control and L1, and 150 for L2. Subplot size was primarily determined to minimise edge effects causing discrepancies between LiDAR and field measurements (60).

### Above-ground biomass disaggregation and net biomass change

The aboveground biomass of individual trees (AGB) was calculated using allometric Eq. 5 of (61): AGB = 0.0559 x (WD x DBH^2^ x H), which incorporates measures of DBH (in cm), height (H, in m), and species wood density (WD, in g cm^-3^). In this equation, the exponent was constrained to one (as opposed to Eq. 4 in (61), which facilitates analytical disaggregation (see below) while keeping good predictive performance. Wood density was extracted from the Global Wood Density Database (62, 63) using the BIOMASS R package (64). Tree height was estimated using a regional diameter–height equation developed for forests of the Guiana Shield (65).

We considered each separate biomass component, based on the approach developed by (29) and applied by (8), in which the stand-level AGB (or AGBplot) estimate, that is, the sum of all individual tree AGB estimates, is analytically disaggregated into three components: the number of trees (N) times the mean individual tree above-ground volume 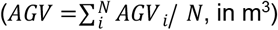 times the mean volume-weighted wood density 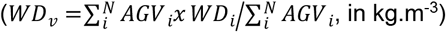. For further details, see (8). After log transformation, we obtain the following additive model:

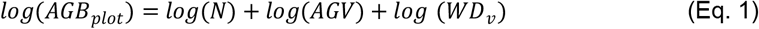

We considered two distinct time intervals—2015 to 2019 and 2019 to 2023—along the forest dynamics trajectory to assess the robustness of our results to different census periods for which LiDAR data were available (see below). In the 2023 inventory, there were 47,317 trees belonging to 590 species. For the time intervals, we computed initial AGB in 2015 and 2019 as baselines, and estimated biomass change in log(AGB) (Δlog(AGB)) over each four-year monitoring interval, i.e., the difference in log(AGB) from the final date to the baseline. According to Eq. 1, Δlog(AGB) is the sum of Δlog(N), Δlog(AGV), and Δlog(WD); henceforth, ΔAGB, ΔNtree, ΔAGV, and ΔWD, respectively.

### Aerial laser scanning - data acquisition and processing

We estimated canopy packing at the subplot scale using airborne laser scanning (ALS) data. ALS data were acquired by the Altoa company, which operated a Riegl LMSQ780 (1064 nm wavelength) onboard a plane flying at ca. 800 m above ground level in 2015 and 2019. The scanning swath angles were 20° and 30° for 2015 and 2019, respectively. The pulse repetition frequency was 400 kHz, for which the aircraft maintained an average ground speed of 180 km h^-1^. The average pulse density was ca. 27 and 42 m^-2^ for 2015 and 2019, respectively. Post-processing of ALS data and point cloud classification into ground, vegetation, or noise was performed using TerraScan (Terrasolid) Version 14. Points classified as ground were used to build a digital terrain model (DTM) at 1-m resolution using LAStools (66). A 1-m resolution canopy height model (CHM) was then computed from the height of the normalised vegetation points by computing the height of the highest return per raster cell.

### Obtaining Plant Area Density (PAD) from AMAPVox

To quantify Plant Area Density (PAD) distributions from ALS data, we estimated canopy light attenuation using AMAPVox. This software tracks every laser pulse through a 3D grid (voxelized space) to the last recorded hit (67). The effective sampling area of each laser pulse (or fraction of pulse in case of multiple hits) is computed from the theoretical beam section (a function of distance to the laser and the divergence of the laser beam) and the remaining beam fraction entering a voxel. In case more than one hit is recorded for a given pulse, then the beam section was equally distributed between the different hits of the pulse. This information is combined with the optical path length of each pulse entering a voxel to compute the local transmittance (the ratio of the sum of energy exiting a voxel to the sum of energy entering the same voxel) or local attenuation per voxel. PAD is then derived from the local attenuation, neglecting within-voxel clumping and assuming isotropic attenuation of light. PAD in each 1 m^3^ voxel thus comprised every structural element of the forest, including non-tree plants such as palms, lianas, and herbs. To evaluate how the different sampling intensities for 2015 and 2019, voxel and subplot size could affect our estimates of PAD, we carried out sensitivity analyses (Appendix S1). For this study, a voxel space of 1 m^3^ was considered large enough to capture the distribution of leaves and trunks, and small enough to represent the forest heterogeneity at the canopy level, thus facilitating the ecological interpretation of PAD spatial arrangement, and a subplot size of 0.25 ha showed acceptable results, striking a balance between spatial resolution and metric stability (Appendix S1).

### Quantifying canopy packing – Shannon Evenness of PAD

To quantify the vertical arrangement of vegetation within a subplot (50m x 50m), we used a canopy index derived from the Shannon Evenness Index (68). This index has previously been used to relate the canopy structural diversity to tropical forest dynamics (26). Here, we computed the Shannon Evenness index, which increases as PAD is distributed more evenly across layers (also called the Vertical Complementarity Index – VCI; (69)). This index is normalised by the total number of layers and is thus, by construction, independent from the total forest height:

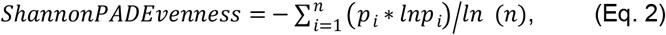

where for every subplot, is the relative PAD value for layer i, computed as the proportion of PAD in that layer relative to the total PAD summed across all layers of the subplot, and is the total number of layers. Here, we calculated this index starting from 5 m aboveground, as lower vegetation layers are poorly sampled and do not allow a reliable estimate of PAD (Appendix S1), to the maximum canopy height of the subplot, using 1-m layer intervals. Since 50m × 50m subplots can include structurally distinct patches (e.g., open gaps and closed canopy) and thus present substantial horizontal heterogeneity in PAD distribution, we also calculated Shannon PAD Evenness at the finer 25m × 25m scale, which could better capture local variation in canopy packing. We then averaged the values across the four 25 m × 25 m quadrats in each 50 m × 50 m subplot. The results were consistent with those obtained using the 50m × 50 m calculation, indicating that our findings are robust to the scale of Shannon index estimation (results not shown).

### Terrain descriptor – Height Above Nearest Drainage (HAND)

To account for the documented impact of local topography on forest dynamics (42, 50), we calculated the Height Above Nearest Drainage (HAND), which relates to downstream drainage networks (70), using the 1-m LiDAR-derived Digital Terrain Model (DTM). High values indicate a greater vertical distance to the nearest drainage, hence lower water availability. The DTM first underwent standard cleaning procedures, including filling small artificial depressions (sinks) smaller than the vertical accuracy of the DTM and smoothing with a moving-window Gaussian filter to remove random errors. We then calculated the HAND index using the ‘whitebox’ R package (71). HAND did not differ among treatments (Fig. S9), indicating comparable topographic conditions across plots.

### Functional traits

We considered several species traits related to resource acquisition, shade tolerance, regeneration strategy, competitive ability, growth and stature. These included leaf area (LA, cm^2^), specific leaf area (SLA, mm^2^/mg), leaf thickness (LT, mm), maximum stem diameter (cm, MaxDBH), and wood specific gravity (WSG, g/cm^3^, hereafter referred to as wood density). Trait data were obtained from previous field campaigns conducted in French Guiana (72). To assess the effects of species traits on Shannon PAD Evenness, we used two types of metrics: univariate (trait-by-trait) and multivariate (all traits) functional dispersion (see below), either weighted by abundance or wood volume (AGV) to account for the dominant role of larger trees in shaping canopy architecture and light interception, which are central to canopy packing. Because FD metrics weighted by abundance exhibited only weak effects on Shannon Evenness of PAD, we present here only the results based on FD metrics weighted by AGV. To quantify functional diversity, a principal coordinates analysis (PCoA) was performed on a species dissimilarity matrix using a square-root correction. We then obtained functional dispersion (FDis), which is estimated as the mean distance of all species to the weighted centroid of the community in the multivariate trait space (73), again weighted by the tree aboveground volume. For univariate analysis, this index corresponded to the weighted mean absolute deviation (MAD, (73)). Only trees identified at the species level and with available trait information were considered, representing ca. 83% of the community volume. Trait-by-trait metrics were also considered because multivariate FD may obscure the potential contributions of individual traits (e.g., (74)). This procedure allowed us to quantify the contributions of individual traits and determine whether trade-offs existed in the magnitude and direction of their effects. Functional diversity indices were calculated in R version 4.3.1 (75) using the package “FD” (73).

### Conceptual model

Here we assumed that canopy packing can increase the number of trees and/or their mean above-ground volume (76), with no causal effect of canopy packing on the community wood density component of AGB, which is instead expected to be related to the sampling effect (8) and will thus not be considered in the following models. The effect of functional diversity on canopy structural diversity (Shannon evenness of PAD, see section below) is thus expected on the number of trees and on above-ground volume (Fig. 1). We evaluated the main conceptual model (Fig. 1) based on the following hypothesised paths: functional dispersion (multivariate and trait-by-trait) calculated at the baseline year of each interval (2015 and 2019) was used as a predictor of Shannon PAD evenness of the same baseline year, which in turn was used as a predictor of biomass gain and loss of the respective intervals (2015–2019 and 2019–2023). We also included the following LiDAR-derived co-variables (also measured at the baseline year): HAND and top-of-canopy height (TCH). HAND was assumed to affect Shannon evenness, ΔAGV and ΔNtree, as water availability was shown to affect canopy structural complexity (53), basal area loss rate (42) and coarse wood productivity (70). Besides, TCH and AGB are positively correlated (77), and AGB has been shown to affect productivity (green soup hypothesis; 78). Finally, as an exact (analytical) relationship, the effect of ΔAGV and ΔNtree on ΔAGB has a known coefficient value of 1 (no estimation is required here due to the analytical solution; Fig. 1).

### Statistical analyses

Piecewise Structural Equation Modelling (pSEM), an extension of the traditional SEM approach, was used to disentangle the effects of trait diversity on biomass gain and biomass loss mediated by canopy packing. In pSEM, the SEM is decomposed into a set of linear models evaluated individually and then combined to make inferences about the entire SEM (79). This approach allows the inclusion of random effects to account for the nested structure of the dataset, thereby accounting for potential pseudo-replication due to subplots nested within larger plots.

We constructed one pSEM for each interval (2015-2019 and 2019-2023) and each treatment (Control, L1, and L2), independently, for the annualised values of the ΔAGB components (ΔAGV and ΔNtree, Mgha^−1^year^−1^). Constructing one model per treatment was necessary to avoid collinearity, as adding a three-level categorical variable substantially increased the Variance Inflation Factor (80). AGB gain and loss were calculated as follows: (i) Recruitment = the sum of the AGB of individuals that recruited into the DBH ≥ 10cm between censuses, divided by the time between measurements. However, for each tree recruited (DBH ≥ 10cm), we subtracted the corresponding AGB associated with a tree of 9.99 cm (i.e., just below the detection limit), to avoid overestimations of the overall productivity in AGB (81, 82). (ii) AGB growth = the sum of the increase in AGB of all individuals with DBH ≥ 10cm that survived between censuses, divided by the time between censuses in years. Biomass gain was then estimated as the sum of biomass recruitment and growth. Biomass loss was estimated as (iii) Mortality = the sum of the AGB of all individuals that died between censuses, divided by the time between measurements. Finally, net change was estimated as the difference between biomass gain and loss.

To evaluate the relative contribution of each predictor to the variance explained in each response variable, we used standardised path coefficients from the piecewise structural equation model.

Although predictors were not scaled prior to model fitting, the standardised estimates provided by the piecewiseSEM package account for differences in variable scales. We computed the relative contribution of each predictor as the absolute value of its standardised coefficient divided by the sum of absolute standardised coefficients for that response, expressed as a percentage. The relative importance of biomass gain (G) and loss (L) to the variation in AGB net change (ΔAGB) was calculated using the ratio of variance and covariance of untransformed variables as described in (83), using the following equations: relative contribution of gains = [var(G + R) − cov(G + R, M)] /var(ΔAGB), and relative contribution of losses = [var(M) − cov(G + R, M)] /var(ΔAGB).

Each model of the piecewise SEM was set up as a linear mixed-effects model (LMM), with a maximum-likelihood fit and the 6.25-ha plots as random intercepts to account for the non-independence of within-plot observations. We used Tukey HSD post hoc pairwise comparisons, performed with the emmeans package (84), to assess differences in Shannon PAD evenness among treatments.

The presence of significant spatial autocorrelation in the model residuals was tested and removed, if present, using the package ‘adespatial’ (85, 86). This step is important to account for all possible residual spatial autocorrelation at different scales (regardless of the random error structure), thereby avoiding spatial statistical artefacts. Once a subset of spatial predictors was selected to remove residual spatial autocorrelation from the initial model, the model was rerun with the additional spatial predictors, yielding unbiased coefficients for the ecological variables of interest.

We also reported two R^2^ values for each response variable: the marginal R^2^ (R^2^m), which represents the variance explained by the fixed effects alone, and the conditional R^2^ (R^2^c), which represents the variance explained by both the fixed and random effects. We evaluated the fit of the entire SEM using Fisher’s C-test; a small C and p>0.05 indicate good model fit. Piecewise SEM was performed in R version 4.5.1 (75), using the packages lme4 (87) and piecewiseSEM (79).

## Acknowledgments

We thank the many colleagues who participated in field and laboratory work for trait data collection in French Guiana. We thank the CIRAD fieldwork team for the tree inventory, and Pascal Petronelli, Giacomo Sellan and Julien Engel for botanical identification. E.R.B. was funded by the European Union’s Horizon 2020 research and innovation programme under the Marie Skłodowska-Curie grant agreement no. 101066995. We thank the Paracou research field station in French Guiana managed and supported by CIRAD, UMR EcoFoG (https://paracou.cirad.fr), which benefits from “Investissement d’Avenir” grants managed by Agence Nationale de la Recherche (AnaEE France ANR-11-INBS-0001; Labex CEBA ANR-10-LABX-25-01).

## Notes

### Competing Interest Statement

The authors have declared no competing interest.

