## Supplementary material for "Trait diversity enhances biomass gains via canopy packing in old-growth but not in disturbed Amazon forests"

PNAS

**This PDF file includes:**

Figs. S1 to S9
Tables S1 to S2
Appendix S1 – Sensitivity analyses

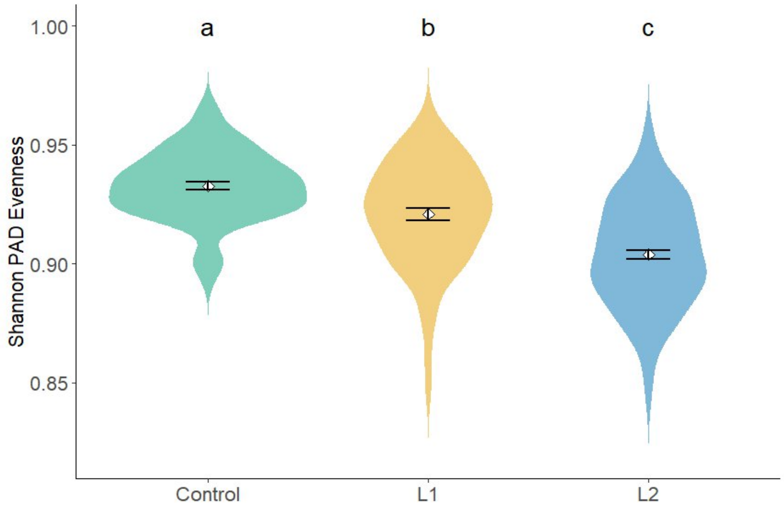


**Fig. S1.**

Violin plots showing the distribution of Shannon' evenness Index of Plant Area Density (PAD) across the logging treatments for 2019. The width of each violin represents the kernel density of values. White diamonds indicate mean values and black error bars represent ±1 standard error of the mean. Different letters (a, b) indicate significant differences in a pairwise comparison of least-squares means (Tukey HSD)

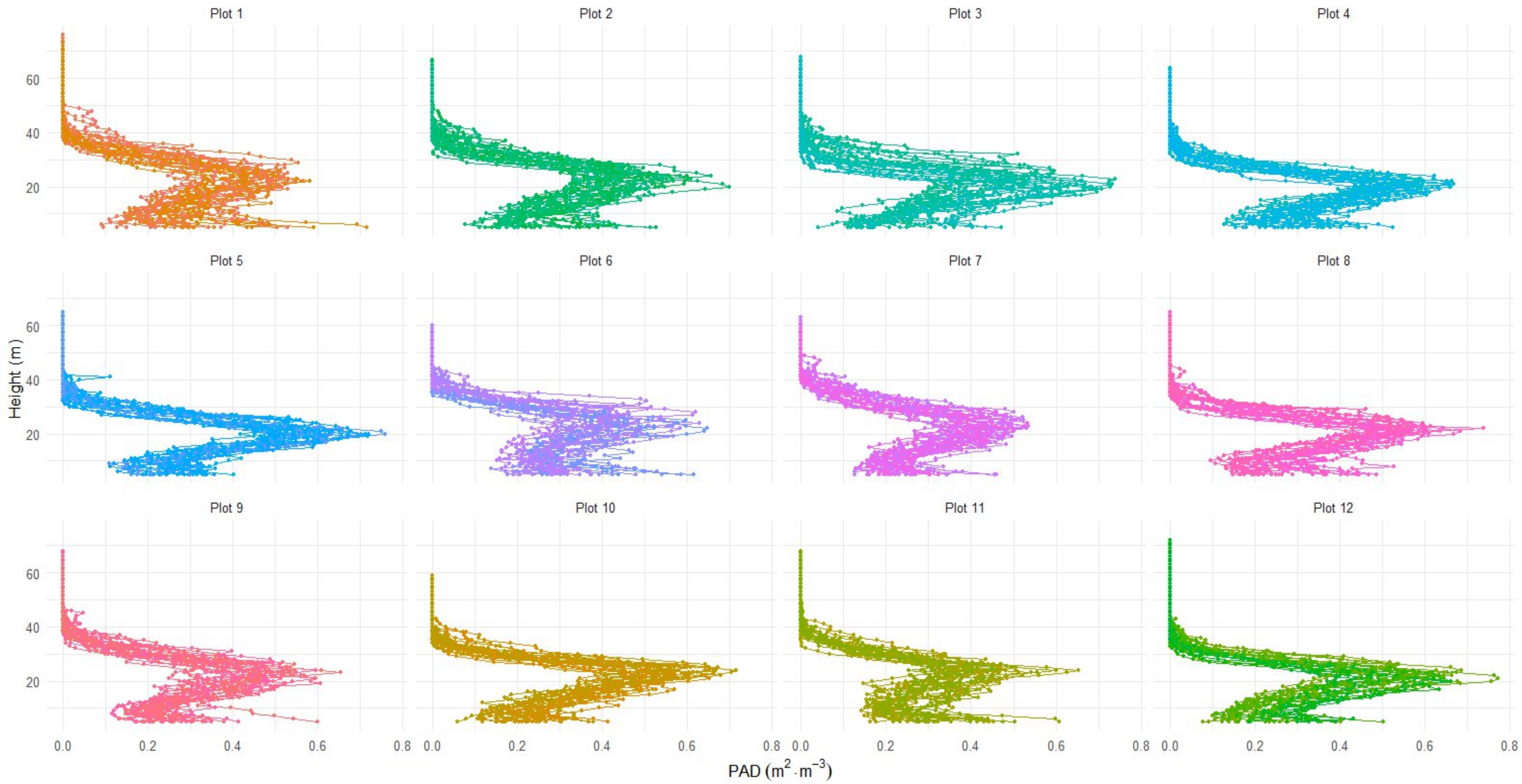


**Fig. S2.**

PAD profiles (vertical distribution curve showing the mean PAD in 1-meter vertical bins) across all forest 0.25-ha subplots illustrating changes in vertical canopy structure in control (Plots 1, 6 and 11), L1(2, 7, 9) and L2 (3, 4, 5, 8, 10, 12) logging treatments. Plant Area Density (PAD) distributions modelled from the 2019 LiDAR data with the AMAPVox software.

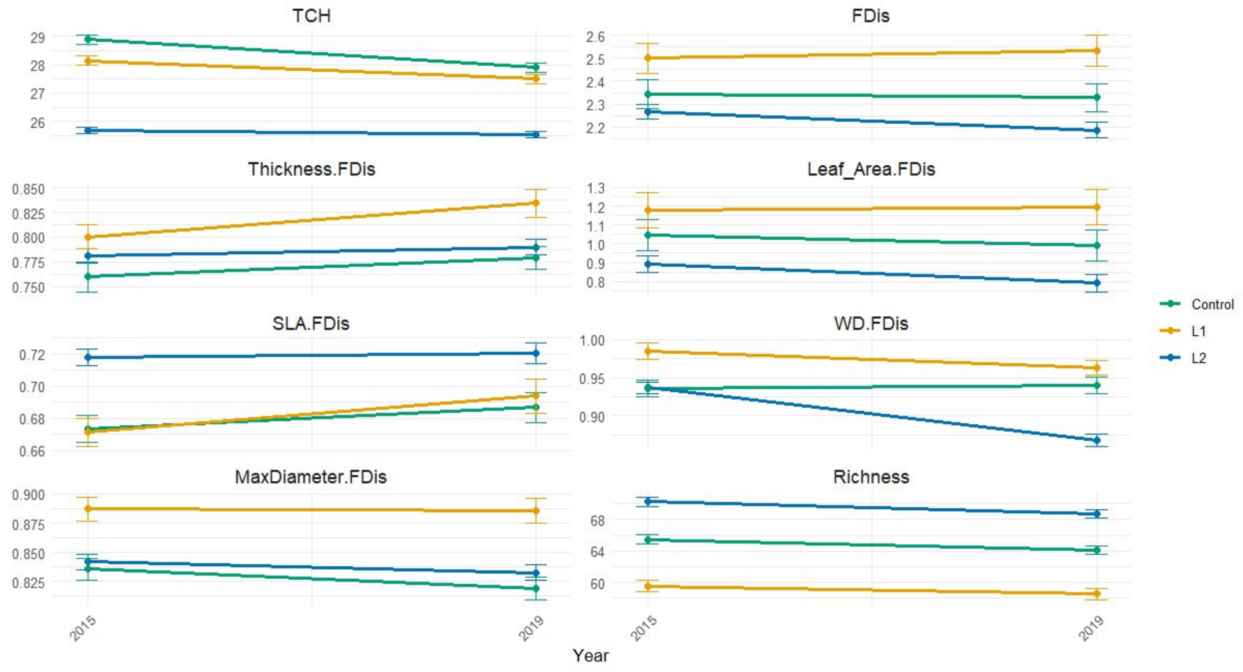


**Fig. S3.**

Effects of disturbance (silvicultural treatments) on multivariate and univariate functional dispersion, richness and TCH over time. Error bars represent 95% confidence interval.

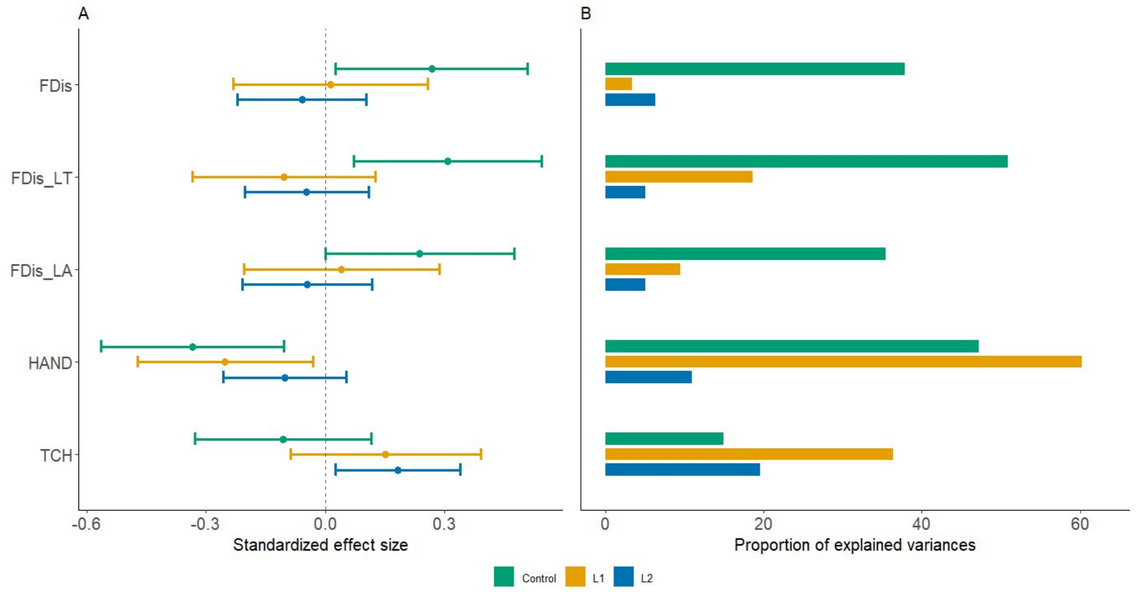


**Fig. S4.**

Standardized effect sizes (A) and proportion of explained variances (B) of the significant predictors of Shannon PAD evenness for 2019. control, L1 and L2. TCH, top-of-canopy-height, HAND, height above nearest drainage, FDis, functional dispersion, LT leaf thickness, LA leaf area.

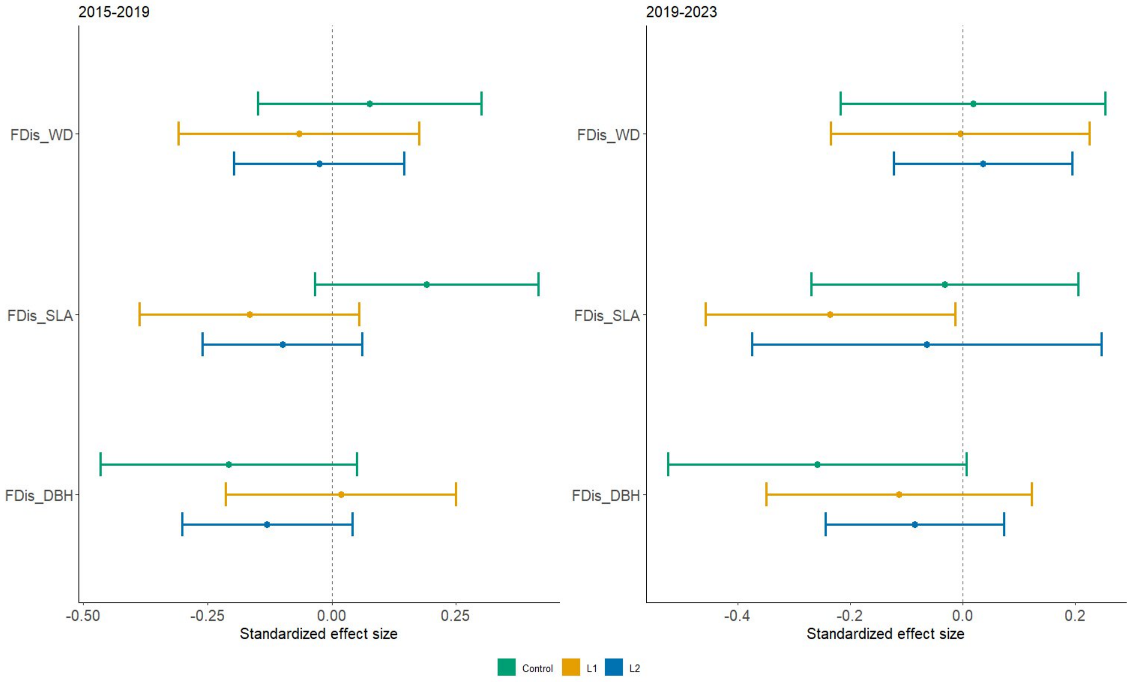


**Fig. S5.**

Standardized effect sizes of univariate functional dispersion on Shannon PAD evenness for both intervals. control, L1 and L2. FDis, functional dispersion, WD wood density, SLA specific leaf area, DBH diameter at breast height.

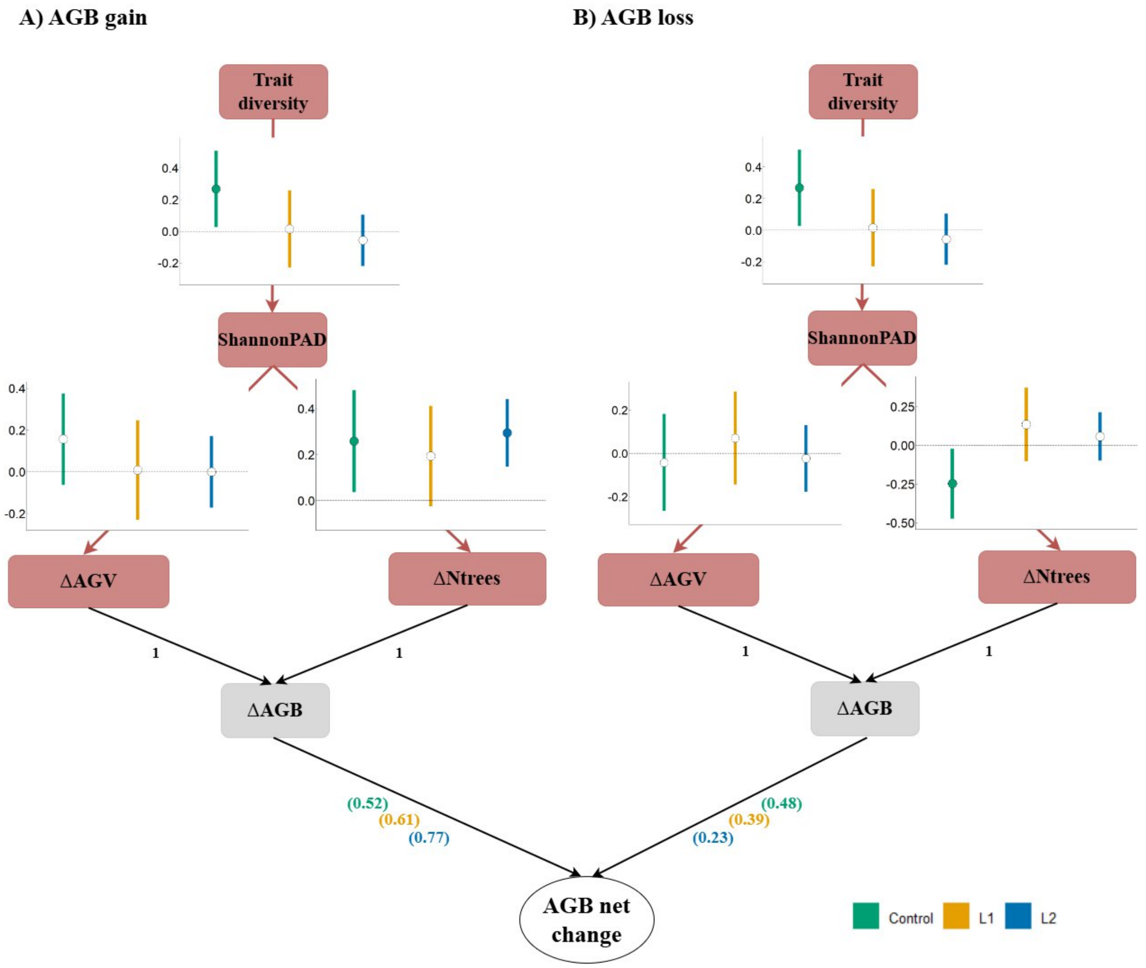


**Fig. S6.**

Results of the structural equation models for the effect of multivariate functional diversity on structural diversity (Shannon PAD) and the components of above-ground biomass gain and loss (ΔAGB) for the first monitoring interval (2019-2023). ΔAGVolume, change in log-transformed mean individual above-ground volume; ΔNtree, change in log transformed number of trees. a) biomass gain, b) biomass loss. The co-variables HAND and CHM are not shown in the SEM for visualization clarity (results shown in Tables S1 and S2).

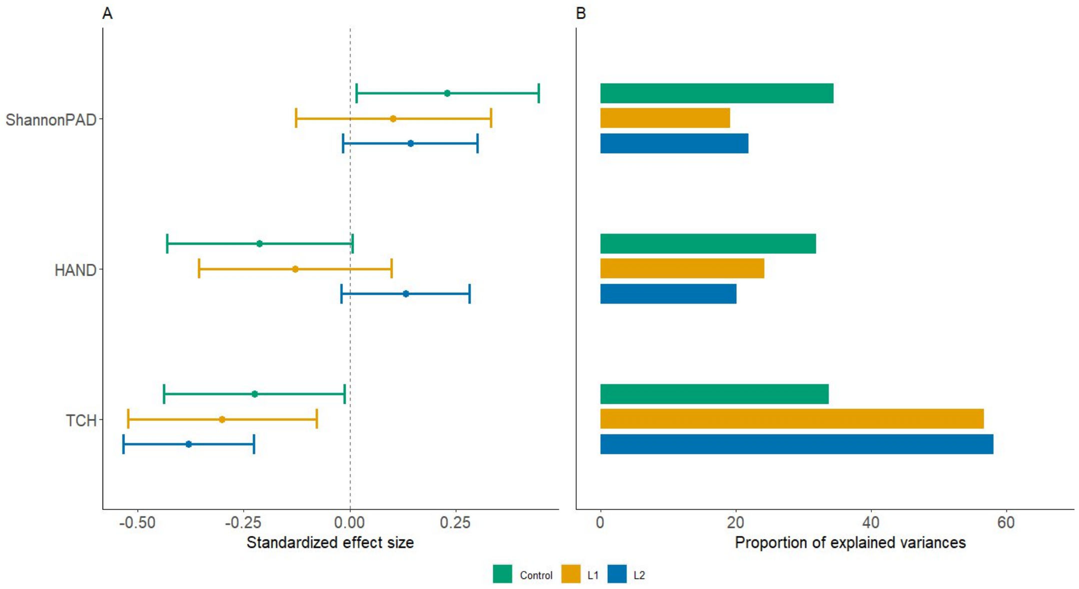


**Fig. S7.**

Standardized effect sizes (A) and proportion of explained variances (B) of the significant predictors of ΔAGB for the 2019-2023 interval for control, L1 and L2 plots. HAND, height above nearest drainage, TCH, top-of-canopy-height.

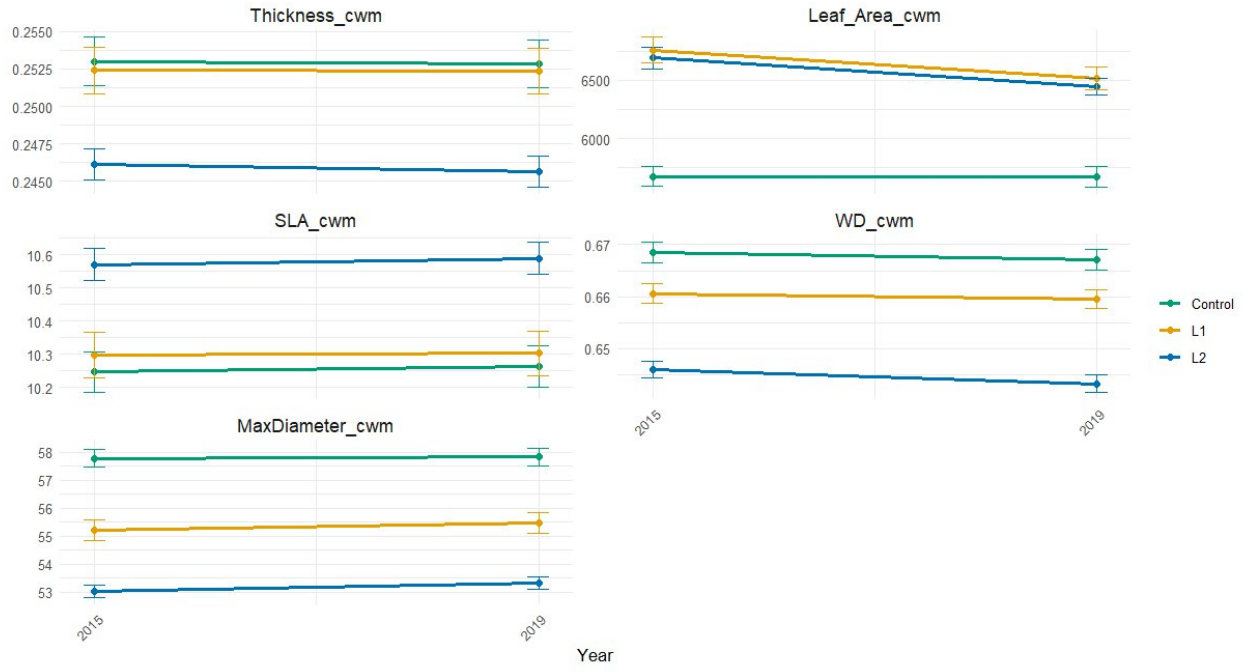


**Fig. S8.**

Effects of disturbance (silvicultural treatments) on community weighted means over time. Error bars represent 95% confidence interval.

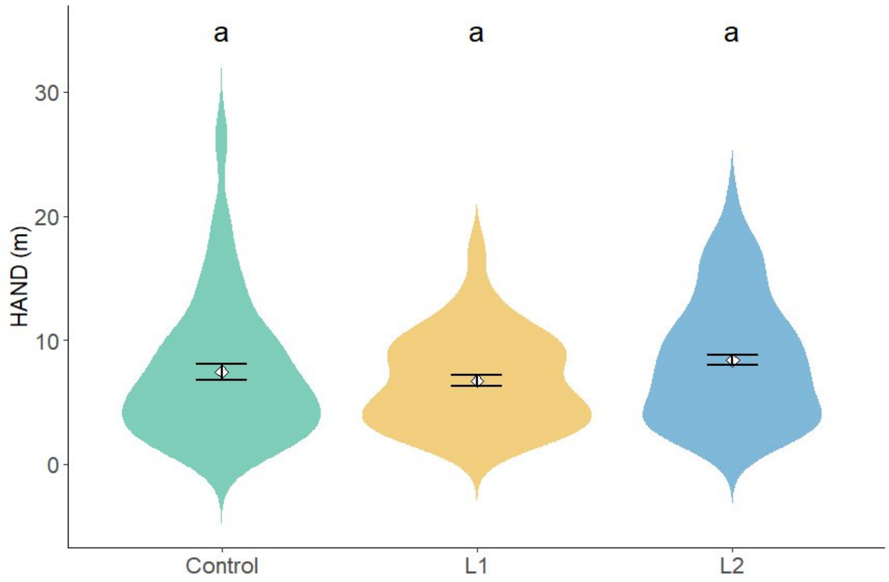


**Fig. S9.**

Violin plots showing the distribution of Height Above Nearest Drainage (HAND) across the logging treatments. The width of each violin represents the kernel density of values. White diamonds indicate mean values and black error bars represent ±1 standard error of the mean. Different letters (a, b) indicate significant differences in a pairwise comparison of least-squares means (Tukey HSD).

| Control | | | | | | | L1 | | | | | L2 | | | | |
|---|---|---|---|---|---|---|---|---|---|---|---|---|---|---|---|---|
| Response | Predictor | Std coef. | S.E. | z-value | *P*-value | Rel. Cont. | Std coef | S.E. | z-value | *P*-value | Rel. Cont. | Std coef. | S.E. | z-value | *P*-value | Rel. Cont. |
| dAGVmeanT | shannonEve_value2015 | 0.23 | 1.74 | 2.02 | 0.05 | 0.71 | 0.10 | 1.13 | 0.85 | 0.40 | 0.20 | 0.01 | 0.63 | 0.17 | 0.86 | 0.05 |
| dAGVmeanT | meanCHM2015 | -0.04 | 0.02 | -0.33 | 0.74 | 0.11 | -0.36 | 0.01 | -3.10 | 0.00 | 0.71 | -0.19 | 0.01 | -2.36 | 0.02 | 0.61 |
| dAGVmeanT | HAND | -0.06 | 0.01 | -0.47 | 0.64 | 0.17 | 0.04 | 0.01 | 0.35 | 0.73 | 0.08 | -0.11 | 0.00 | -1.33 | 0.19 | 0.34 |
| dNtree2019 | shannonEve_value2015 | 0.37 | 3.73 | 3.15 | 0.00 | 0.48 | 0.19 | 2.66 | 1.56 | 0.12 | 0.29 | 0.23 | 1.51 | 2.96 | 0.00 | 0.24 |
| dNtree2019 | meanCHM2015 | -0.22 | 0.03 | -1.89 | 0.06 | 0.28 | -0.25 | 0.03 | -2.11 | 0.04 | 0.37 | -0.38 | 0.02 | -4.80 | 0.00 | 0.41 |
| dNtree2019 | HAND | 0.19 | 0.01 | 1.57 | 0.12 | 0.24 | 0.22 | 0.02 | 1.86 | 0.07 | 0.34 | 0.08 | 0.01 | 1.01 | 0.32 | 0.08 |
| shannonEve_value2015 | FDis | 0.23 | 0.00 | 2.02 | 0.05 | 0.33 | 0.10 | 0.00 | 0.80 | 0.43 | 0.22 | -0.01 | 0.00 | -0.07 | 0.94 | 0.01 |
| shannonEve_value2015 | HAND | -0.41 | 0.00 | -3.42 | 0.00 | 0.58 | -0.26 | 0.00 | -2.46 | 0.02 | 0.58 | -0.13 | 0.00 | -1.59 | 0.11 | 0.15 |
| shannonEve_value2015 | meanCHM2015 | -0.06 | 0.00 | -0.51 | 0.61 | 0.08 | 0.09 | 0.00 | 0.79 | 0.43 | 0.20 | 0.18 | 0.00 | 2.27 | 0.02 | 0.22 |
| ~~dNtree2019 | ~~dAGVmeanT | 0.27 | - | 2.35 | 0.01 | 1.00 | 0.43 | - | 4.02 | 0.00 | 1.00 | 0.20 | - | 2.41 | 0.01 | 1.00 |
| shannonEve_value2015 | MEMSH1 | | | | | | | | | | | -0.28 | 0.00 | -3.73 | 0.00 | 0.33 |
| shannonEve_value2015 | MEMSH2 | | | | | | | | | | | -0.26 | 0.00 | -3.44 | 0.00 | 0.30 |
| dAGBT | shannonEve_value2015 | 0.32 | 2.18 | 2.89 | 0.01 | 0.71 | 0.15 | 1.54 | 1.29 | 0.20 | 0.24 | 0.12 | 0.81 | 1.51 | 0.13 | 0.27 |
| dAGBT | meanCHM2015 | -0.11 | 0.02 | -1.04 | 0.30 | 0.25 | -0.36 | 0.02 | -3.19 | 0.00 | 0.57 | -0.30 | 0.01 | -3.78 | 0.00 | 0.69 |
| dAGBT | HAND | 0.02 | 0.01 | 0.19 | 0.85 | 0.05 | 0.12 | 0.01 | 1.06 | 0.29 | 0.19 | -0.02 | 0.00 | -0.22 | 0.83 | 0.04 |
| $R^2$ m dAGVmean | | 0.07 | | | | | 0.12 | | | | | 0.05 | | | | |
| $R^2$ c dAGVmean | | 0.21 | | | | | 0.12 | | | | | 0.05 | | | | |
| $R^2$ m dNtree | | 0.17 | | | | | 0.10 | | | | | 0.19 | | | | |
| $R^2$ c dNtree | | 0.17 | | | | | 0.10 | | | | | 0.19 | | | | |
| $R^2$ m ShannonPAD | | 0.17 | | | | | 0.10 | | | | | 0.17 | | | | |
| $R^2$ c ShannonPAD | | 0.17 | | | | | 0.23 | | | | | 0.21 | | | | |
| $R^2$ m dAGB | | 0.12 | | | | | 0.14 | | | | | 0.10 | | | | |
| $R^2$ c dAGB | | 0.25 | | | | | 0.14 | | | | | 0.11 | | | | |
| *Model goodness of fit* | | | | | | | | | | | | | | | | |
| $X^2$ | | 1.10 | | | | | 3.89 | | | | | 5.31 | | | | |
| p-value $X^2$ | | 0.58 | | | | | 0.14 | | | | | 0.38 | | | | |
| Fisher's C | | 2.98 | | | | | 7.62 | | | | | 10.69 | | | | |
| p-value Fisher's C | | 0.56 | | | | | 0.11 | | | | | 0.38 | | | | |

**Table S1.**

Results of the piecewise structural equation models for aboveground biomass gain (2015-2019). Results show direct and indirect effect, $R^2$ and model goodness of fit.

| | | Control | | | | | L1 | | | | | L2 | | | | |
|---|---|---|---|---|---|---|---|---|---|---|---|---|---|---|---|---|
| Response | Predictor | Std coef. | S.E. | z-value | *P*-value | Rel. Cont. | Std coef. | S.E. | z-value | *P*-value | Rel. Cont. | Std coef. | S.E. | z-value | *P*-value | Rel. Cont. |
| dAGVmeanT | shannonEve_value2019 | 0.16 | 1.89 | 1.42 | 0.16 | 0.27 | 0.01 | 1.18 | 0.07 | 0.94 | 0.01 | 0.00 | 0.91 | -0.02 | 0.99 | 0.01 |
| dAGVmeanT | meanCHM2019 | -0.22 | 0.01 | -2.01 | 0.05 | 0.39 | -0.18 | 0.01 | -1.56 | 0.12 | -0.18 | -0.23 | 0.01 | -2.72 | 0.01 | 0.87 |
| dAGVmeanT | HAND | -0.19 | 0.01 | -1.72 | 0.09 | 0.34 | -0.03 | 0.01 | -0.25 | 0.80 | -0.03 | -0.03 | 0.00 | -0.40 | 0.69 | 0.13 |
| dNtree2023 | shannonEve_value2019 | 0.26 | 1.13 | 2.28 | 0.03 | 0.44 | 0.19 | 0.74 | 1.74 | 0.09 | 0.19 | 0.29 | 0.49 | 3.93 | 0.00 | 0.31 |
| dNtree2023 | meanCHM2019 | -0.16 | 0.01 | -1.43 | 0.16 | 0.27 | -0.34 | 0.01 | -3.17 | 0.00 | -0.34 | -0.38 | 0.01 | -5.21 | 0.00 | 0.40 |
| dNtree2023 | HAND | -0.17 | 0.00 | -1.52 | 0.13 | 0.30 | -0.22 | 0.00 | -1.96 | 0.05 | -0.22 | 0.29 | 0.00 | 3.97 | 0.00 | 0.30 |
| shannonEve_value2019 | FDis | 0.27 | 0.00 | 2.23 | 0.03 | 0.39 | 0.00 | 0.00 | -0.01 | 0.99 | 0.00 | -0.04 | 0.00 | -0.43 | 0.67 | 0.17 |
| shannonEve_value2019 | HAND | -0.33 | 0.00 | -2.83 | 0.01 | 0.47 | -0.25 | 0.00 | -2.24 | 0.03 | -0.25 | -0.05 | 0.00 | -0.60 | 0.55 | 0.23 |
| shannonEve_value2019 | meanCHM2019 | -0.10 | 0.00 | -0.92 | 0.36 | 0.15 | 0.16 | 0.00 | 1.30 | 0.20 | 0.16 | 0.14 | 0.00 | 1.61 | 0.11 | 0.61 |
| ~~dNtree2023 | ~~dAGVmeanT | 0.32 | - | 2.89 | 0.00 | 1.00 | 0.32 | - | 2.88 | 0.00 | 0.32 | 0.08 | - | 0.97 | 0.17 | 1.00 |
| dAGBT | shannonEve_value2019 | 0.23 | 2.51 | 2.10 | 0.04 | 0.34 | 0.10 | 1.58 | 0.87 | 0.39 | 0.19 | 0.14 | 1.07 | 1.76 | 0.08 | 0.22 |
| dAGBT | meanCHM2019 | -0.23 | 0.02 | -2.07 | 0.04 | 0.34 | -0.30 | 0.02 | -2.66 | 0.01 | 0.57 | -0.38 | 0.01 | -4.84 | 0.00 | 0.58 |
| dAGBT | HAND | -0.21 | 0.01 | -1.91 | 0.06 | 0.32 | -0.13 | 0.01 | -1.11 | 0.27 | 0.24 | 0.13 | 0.00 | 1.70 | 0.09 | 0.20 |
| $R^2$ m dAGVmean | | 0.15 | | | | | 0.04 | | | | | 0.05 | | | | |
| $R^2$ c dAGVmean | | 0.23 | | | | | 0.04 | | | | | 0.05 | | | | |
| $R^2$ m dNtree | | 0.17 | | | | | 0.20 | | | | | 0.27 | | | | |
| $R^2$ c dNtree | | 0.17 | | | | | 0.20 | | | | | 0.29 | | | | |
| $R^2$ m ShannonPAD | | 0.15 | | | | | 0.09 | | | | | 0.02 | | | | |
| $R^2$ c ShannonPAD | | 0.18 | | | | | 0.11 | | | | | 0.02 | | | | |
| $R^2$ m dAGB | | 0.21 | | | | | 0.12 | | | | | 0.16 | | | | |
| $R^2$ c dAGB | | 0.24 | | | | | 0.12 | | | | | 0.16 | | | | |
| *Model goodness of fit* | | | | | | | | | | | | | | | | |
| $X^2$ | | 1.45 | | | | | 0.24 | | | | | 5.68 | | | | |
| p-value $X^2$ | | 0.49 | | | | | 0.89 | | | | | 0.06 | | | | |
| Fisher's C | | 3.53 | | | | | 1.01 | | | | | 9.37 | | | | |
| p-value Fisher's C | | 0.47 | | | | | 0.91 | | | | | 0.05 | | | | |

**Table S2.**

Results of the piecewise structural equation models for aboveground biomass gain (2019-2023). Results show direct and indirect effect, $R^2$ and model goodness of fit.

**Appendix S1 – Sensitivity analyses**

1. Data

Using AMAPVox we created 3D voxel grids based on LIDAR point cloud data for each Paracou plot.

Thinning Levels Tested for Paracou Plot1:

- Original: 161.95 pulses/m²
- Level 1: 34.82 pulses/m² ~ 35
- Level 2: 9.95 pulses/m² ~ 10
- Level 3: 4.98 pulses/m² ~ 5

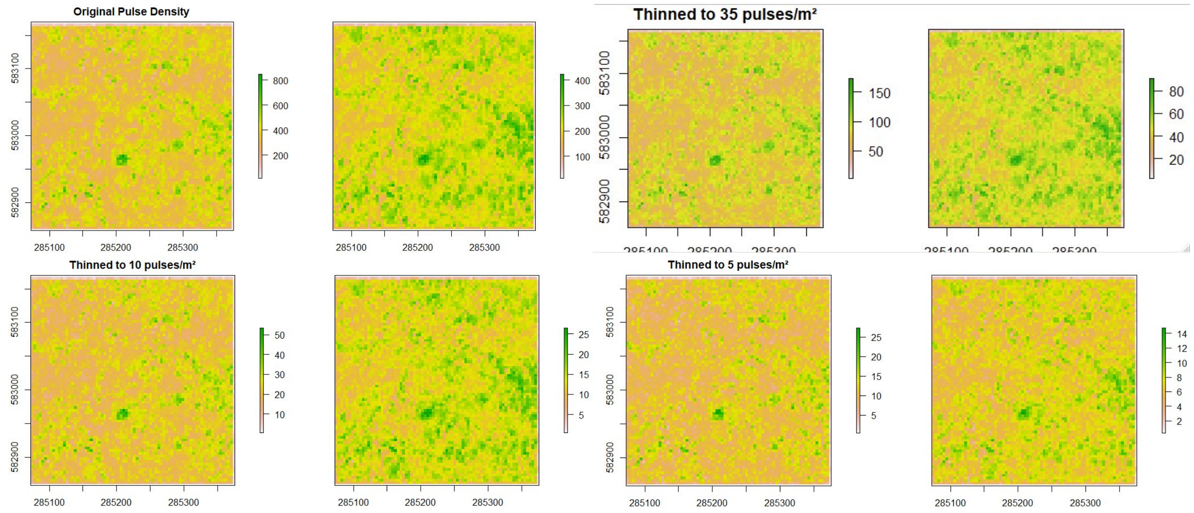


2. Why 1m voxel? - Voxel size testing

Two voxel resolutions were tested: 1m × 1m × 1m voxels and 2m × 2m × 2m voxels. (SubPlot=0.5ha, grid3*3, Cap attPPL=1.5)

From the 9-subplot comparison, **attPPL with 1m voxel emerges as the most stable metric across all pulse density**.

1m voxels: Show remarkable stability across pulse density, with profiles remaining closely aligned from original to highly thinned datasets; 2m voxels (label: xxxxx_2.vox): Demonstrate greater divergence between different pulse densities, with more pronounced separation between profile lines.

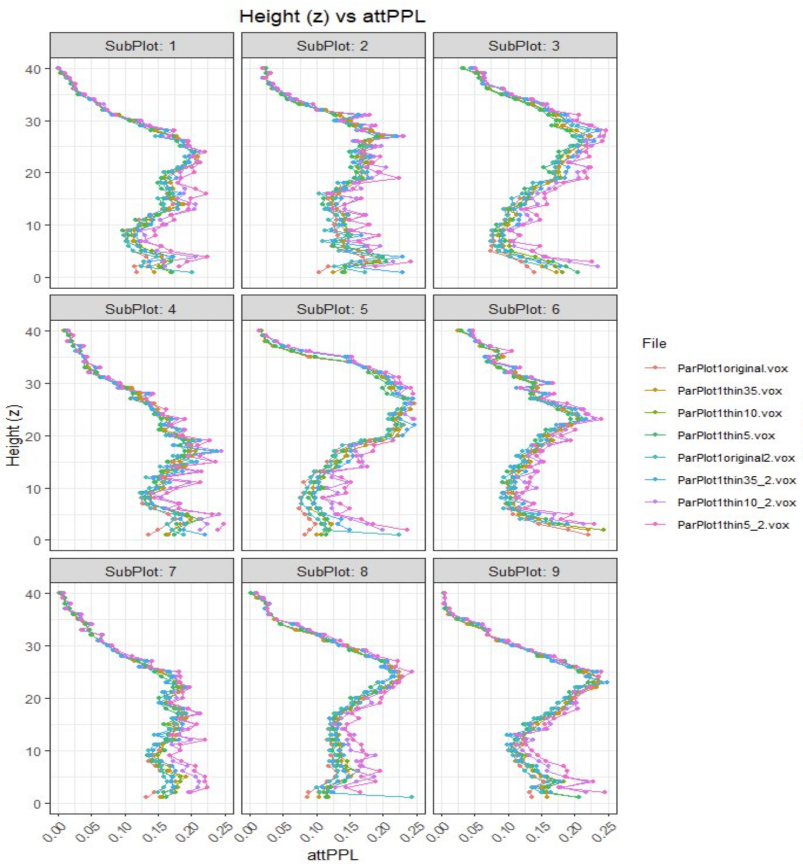

Height (z) vs attPPL
SubPlot: 1
SubPlot: 2
SubPlot: 3
SubPlot: 4
SubPlot: 5
SubPlot: 6
SubPlot: 7
SubPlot: 8
SubPlot: 9
Height (z)
attPPL
File
ParPlot1original.vox
ParPlot1thin35.vox
ParPlot1thin10.vox
ParPlot1thin5.vox
ParPlot1original2.vox
ParPlot1thin35_2.vox
ParPlot1thin10_2.vox
ParPlot1thin5_2.vox


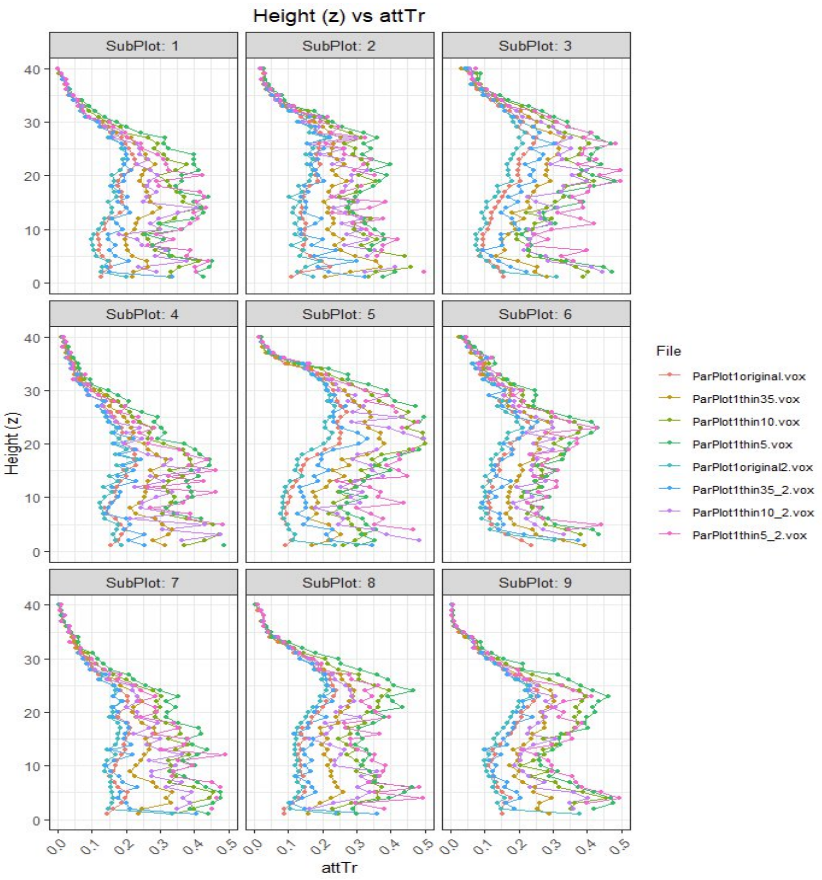

Height (z) vs attTr
SubPlot: 1
SubPlot: 2
SubPlot: 3
SubPlot: 4
SubPlot: 5
SubPlot: 6
SubPlot: 7
SubPlot: 8
SubPlot: 9
Height (z)
attTr
File
ParPlot1original.vox
ParPlot1thin35.vox
ParPlot1thin10.vox
ParPlot1thin5.vox
ParPlot1original2.vox
ParPlot1thin35_2.vox
ParPlot1thin10_2.vox
ParPlot1thin5_2.vox

3. Why 3*3 resolution? - Spatial resolution testing

Five different spatial resolutions were tested by subdividing each plot into varying numbers of subplots (using optimal parameters: Voxel size=1m, Cap value_attPPL=1.5).

| Resolution | Subplot area | Edge length |
|---|---|---|
| 2-2 | 1.5ha | 114~115m |
| 3-3 | 0.58~0.59ha | 76~78m |
| 4-4 | 0.33ha | 57m |
| 7-7 | 0.1ha | 33m |
| 10-10 | 0.05ha | 23m |

The 3×3 grid resolution (0.58-0.59ha subplots) demonstrates the most stable and consistent difference patterns. Finer resolutions (7×7 and 10×10) show increased variability. Coarser resolutions (2×2) show acceptable stability but provide less spatial detail for heterogeneity analysis. The 3×3 resolution offers the optimal balance between spatial resolution and metric stability, but also provides less spatial detail for heterogeneity analysis. The 4×4 resolution (~ 57m x 57m, slightly larger than the 50m x 50m subplots) represents the best compromise of spatial resolution and metric stability (see below) and was thus the choice of the subplot size in this study.

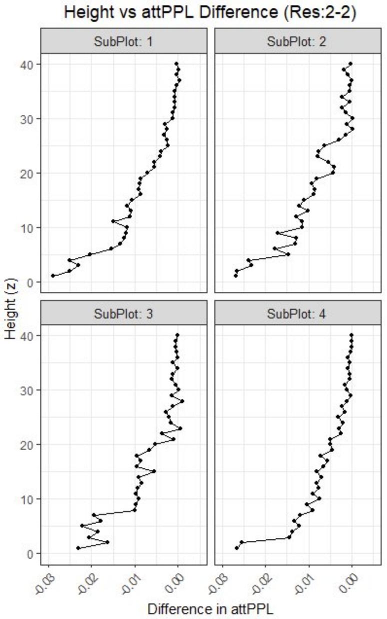
Height vs attPPL Difference (Res:2-2)
SubPlot: 1
SubPlot: 2
SubPlot: 3
SubPlot: 4
Height (z)
Difference in attPPL

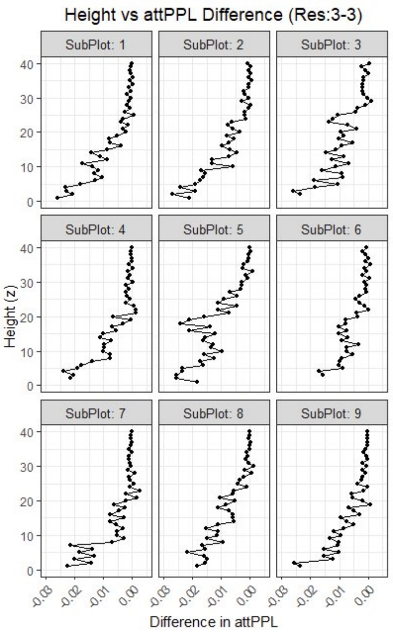
Height vs attPPL Difference (Res:3-3)
SubPlot: 1
SubPlot: 2
SubPlot: 3
SubPlot: 4
SubPlot: 5
SubPlot: 6
SubPlot: 7
SubPlot: 8
SubPlot: 9
Height (z)
Difference in attPPL

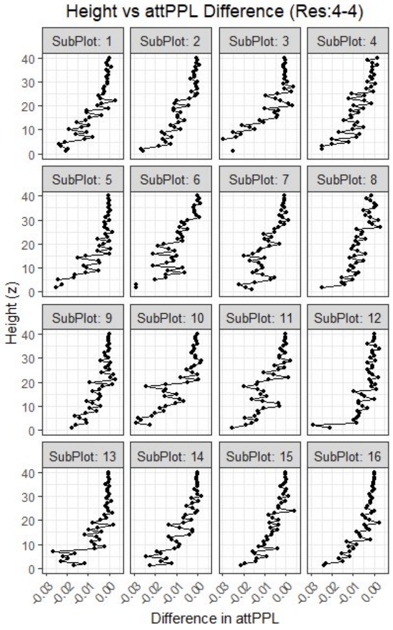
Height vs attPPL Difference (Res:4-4)
SubPlot: 1
SubPlot: 2
SubPlot: 3
SubPlot: 4
SubPlot: 5
SubPlot: 6
SubPlot: 7
SubPlot: 8
SubPlot: 9
SubPlot: 10
SubPlot: 11
SubPlot: 12
SubPlot: 13
SubPlot: 14
SubPlot: 15
SubPlot: 16
Height (z)
Difference in attPPL

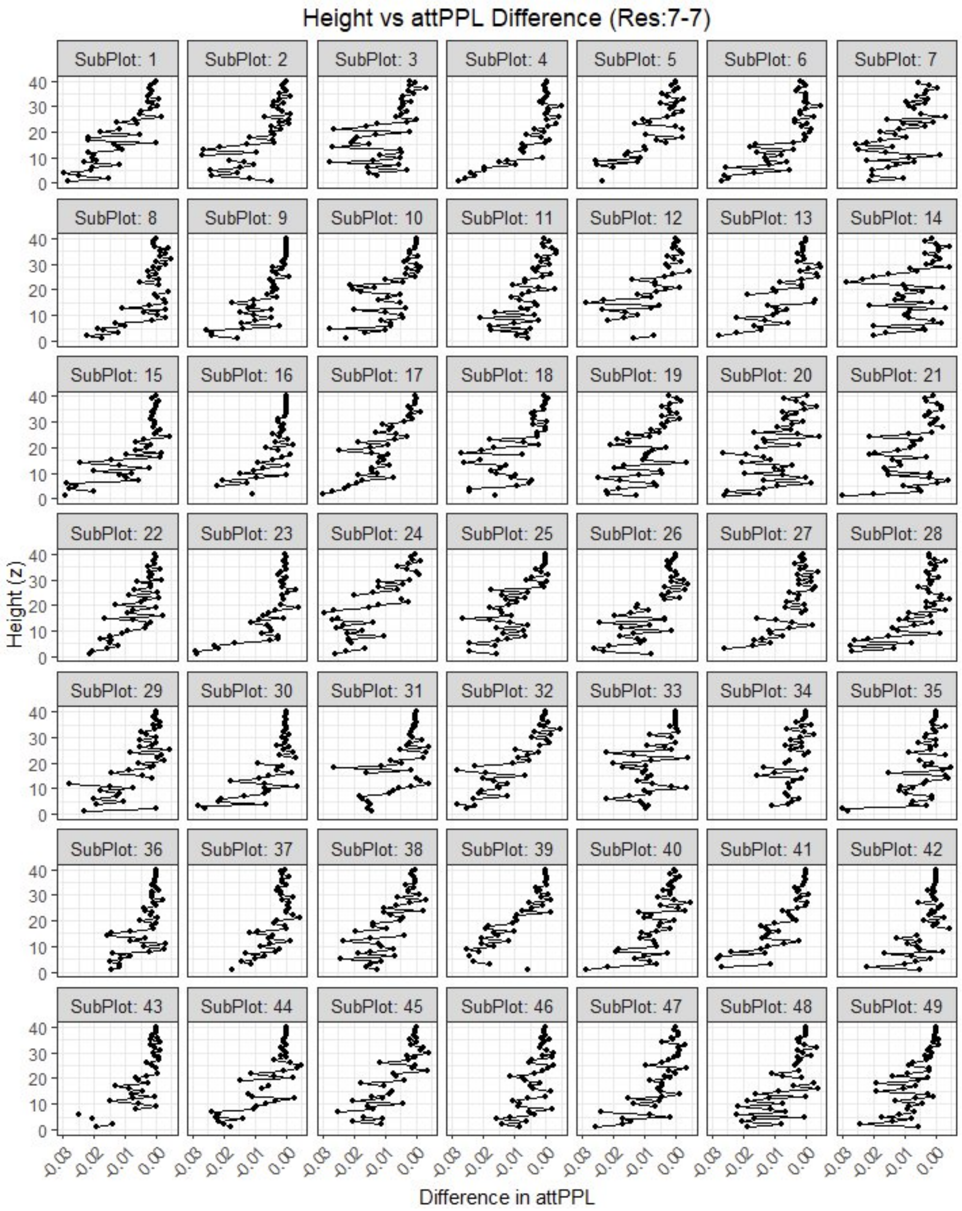

Height vs attPPL Difference (Res:7-7)
SubPlot: 1
SubPlot: 2
SubPlot: 3
SubPlot: 4
SubPlot: 5
SubPlot: 6
SubPlot: 7
SubPlot: 8
SubPlot: 9
SubPlot: 10
SubPlot: 11
SubPlot: 12
SubPlot: 13
SubPlot: 14
SubPlot: 15
SubPlot: 16
SubPlot: 17
SubPlot: 18
SubPlot: 19
SubPlot: 20
SubPlot: 21
SubPlot: 22
SubPlot: 23
SubPlot: 24
SubPlot: 25
SubPlot: 26
SubPlot: 27
SubPlot: 28
SubPlot: 29
SubPlot: 30
SubPlot: 31
SubPlot: 32
SubPlot: 33
SubPlot: 34
SubPlot: 35
SubPlot: 36
SubPlot: 37
SubPlot: 38
SubPlot: 39
SubPlot: 40
SubPlot: 41
SubPlot: 42
SubPlot: 43
SubPlot: 44
SubPlot: 45
SubPlot: 46
SubPlot: 47
SubPlot: 48
SubPlot: 49
Height (z)
0
10
20
30
40
-0.03
-0.02
-0.01
0.00
Difference in attPPL
